# An integrated platform for simultaneous wide-field voltage/calcium imaging and fMRI (EPI & ZTE) reveals neuronal infraslow dynamics underlying functional connectivity

**DOI:** 10.64898/2026.01.26.701889

**Authors:** Wen-Ju Pan, Lauren Daley, Harrison Watters, Lisa Meyer-Baese, Kaudinya Gopinath, Dieter Jaeger, Shella D. Keilholz

## Abstract

Wide-field optical imaging acquired simultaneously with functional MRI (fMRI) has the ability to provide unprecedented insight into the neural origins of time-varying whole-brain activity. Simultaneously linking cellular-scale activity to whole-brain fMRI remains challenging due to optical access, RF coil placement, and transmission constraints in the MRI environment. We present an integrated platform that combines a long-distance tube-lens optical path (>98% transmission), a chronically stable optically-fused cranial window, and a subject-conformal RF surface coil compatible with both echo planar imaging (EPI) and zero-echo-time (ZTE) fMRI. The system supports concurrent wide-field imaging of genetically encoded voltage or calcium indicators concurrently with intrinsic hemoglobin signals. In individual mice, wide-field optical and fMRI measures yield concordant functional connectivity, and cross-modal timing analyses demonstrate that neuronal infraslow dynamics (<0.1 Hz) underlie the majority of fMRI connectivity, after removing hemodynamic crosstalk. The platform’s sensitivity, chronic stability, and sequence flexibility broaden access to cellular-to-whole-brain investigations across basic and translational neuroimaging.

## Main

Noninvasive mapping of brain-wide networks remains a central goal in systems neuroscience. Functional MRI (fMRI) uniquely provides whole-brain coverage in humans and animal models, and extensive work has linked BOLD fluctuations to underlying neuronal activity and neurovascular coupling^1–6^. However, most direct multimodal studies in animals have been spatially restricted, limiting network-level interpretation^4,7,8^. Wide-field optical imaging (WOI) bridges this gap by enabling cortex-wide readouts of genetically targeted calcium or voltage signals, providing cellularly-informed measurements at mesoscopic scales. Simultaneous WOI–fMRI enables time-locked comparison of cellular signals to whole-brain hemodynamics, but prior systems have faced constraints, including calcium-only compatibility, low optical transmission from fiber bundles, and interference between optical windows and RF coils^7^. We address these limitations with a platform that: (i) uses a long-distance tube-lens to position a scientific camera outside the magnet room with >98% transmission; (ii) employs a chronically stable optically cured thin-bone cranial window; (iii) integrates a subject-conformal RF surface coil; and (iv) provides flexible multi-channel illumination for GEVI/GECI and intrinsic signals. The platform supports both EPI and ZTE fMRI, yields concordant single-animal functional connectivity across modalities, and enables separation of neuronal and hemodynamic contributions in low frequency bands—implicating neuronal infraslow dynamics in network organization^2,9^.

### Efficient MRI-compatible wide-field optical imaging system

We adapted a benchtop WOI system for MRI compatibility by routing the reflected and emitted light to a camera outside the magnet room via a customized long-distance tube-lens (Figure 1). The optical train (objective–relay–camera lenses, 400–1000 nm coatings) maintained >98% transmission across ~4 m. The objective was positioned ~80 cm from the cortex to cover an ~10 mm field of view. A 3-to-2 fiber bundle provided multi-channel illumination. We interleaved GEVI or GECI fluorescence (blue excitation) with intrinsic green reflectance (525/50 nm), the latter near an isosbestic point to estimate HbT. HbT was used both as a physiological readout and to regress hemodynamic absorption from fluorescence to minimize crosstalk^16,18^.

**Fig. 1.**
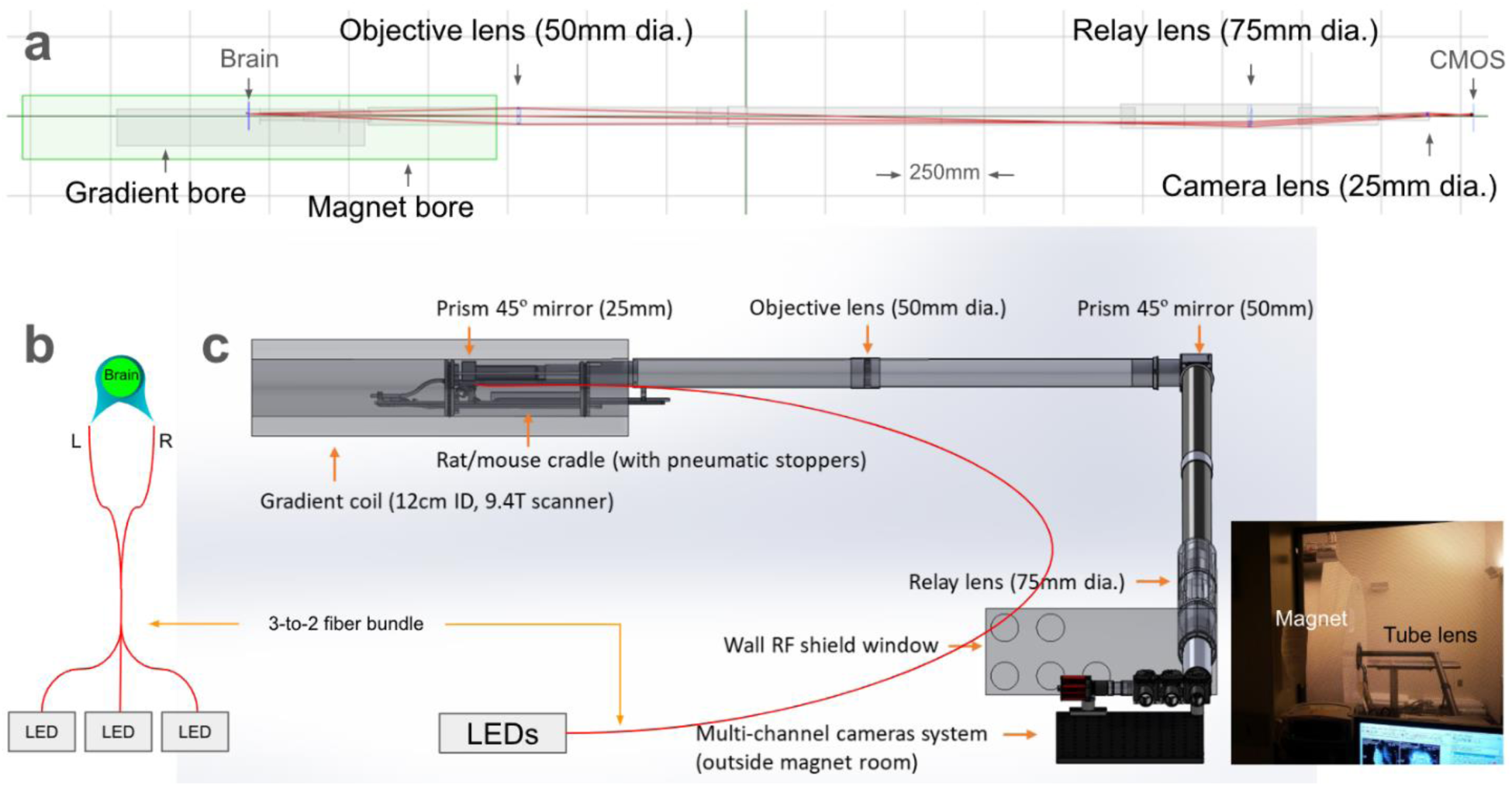
WOI/ fMRI integration with long tube-lens. (a) Schematic of the optical path (brain-objective– relay–camera lenses-CMOS; ~4 m total), with mirrors to route light from the magnet isocenter to a scientific camera outside the magnet room. (b) Multi-channel, bilateral fiber-optic illumination allows interleaved GEVI/GECI fluorescence and intrinsic reflectance (HbT). (c) Custom cradle with head holder accommodates the optically fused cranial window and subject-conformal RF coil (Figure 2); pneumatic stoppers reduce vibration.

### Transparent cranial window and efficient RF surface coil combination

We implemented a chronically stable, optically cured thin-bone cranial window tailored for simultaneous WOI and fMRI. An 8–10 mm circular area over the dorsal cortex was thinned to the inner table; bone channels were removed to improve clarity. A glass coverslip was bonded to the thinned bone using a refractive-index-matched optical adhesive (UV-cured), producing a flat, glare-minimized surface with mechanical reinforcement. A lightweight 3D-printed window frame and removable head-holding bar enabled secure fixation without ear bars, facilitating awake or anesthetized imaging. The frame fit within an oval, curved RF surface coil (10 × 15 mm ID) positioned to maximize whole-brain coverage and B_1_ orientation relative to B_0_.

### Zero- echo time (ZTE) fMRI for quiet and distortion-free scanning

The vast majority of fMRI studies utilize an echo-planar imaging (EPI) sequence, but these sequences are vulnerable to susceptibility artifacts and signal loss, and produce very loud acoustic noise. Zero- echo time (ZTE) fMRI is a radial imaging sequence recently adopted for fMRI that offers several advantages over traditional fMRI methods, including a relatively quiet scanning process, reduced susceptibility to artifacts, and the potential for capturing signals from short-T2 tissues and lower brain regions that are typically missing in EPI due to signal dropout ^10^. ZTE is particularly well-suited for multimodal experiments, as its unique pulse sequence design can reduce the impact of motion and susceptibility artifacts due to surgery, a key enabling feature for future experiments (e.g., with electrodes, deep stimulation, awake studies) where EPI quality is typically low ^11,12^. To show the flexibility of our WOI/fMRI system, we demonstrate high quality fMRI with both pulse sequences, Figure S3. The position of the RF coil in tight proximity to the brain facilitates high power delivery efficiently when a very brief pulse is employed in the ZTE sequence.

## Results

To evaluate the suitability of our fMRI/WOI system for a range of experiments, image quality and functional connectivity (FC) were examined for neuronal fluorescence signals from calcium or voltage indicators, reflectance signals of total hemoglobin absorption (HbT), and EPI/ZTE fMRI signals acquired simultaneously from lightly anesthetized mice ^13^. All data were preprocessed and registered to the Allen atlas, and the spatial and temporal consistency of the fMRI and WOI results were examined.

### A Novel Cranial Window and RF Coil Design Enables High-Quality Dual-Modal Imaging

We first evaluated the image quality control for simultaneously-acquired WOI and fMRI (EPI or ZTE). Qualitatively, images from both modalities acquired simultaneously appeared similar to images acquired separately. The WOI qualitybenefited from the new cranial window design, demonstrated in Figure 2, which resulted in improved transparency and long-lasting durability. The designed cranial implantation also benefited fMRI on optimizing a removable surface coil positioning with minimum coil-brain distance and reproducibility, which resulted in a more sensitive signal detection and higher SNR than that obtained previously using a conventional surface coil setup, reported in ISMRM ^14^. A direct comparison of fMRI with ZTE and EPI from our pilot studies is shown in supplementary Figure S1, with both fMRI sequences demonstrating the ability to detect functional networks in a single scan from a single mouse. Advantages and disadvantages of the conventional fMRI and simultaneous fMRI/WOI approaches are summarized in supplementary table S1.

**Fig. 2.**
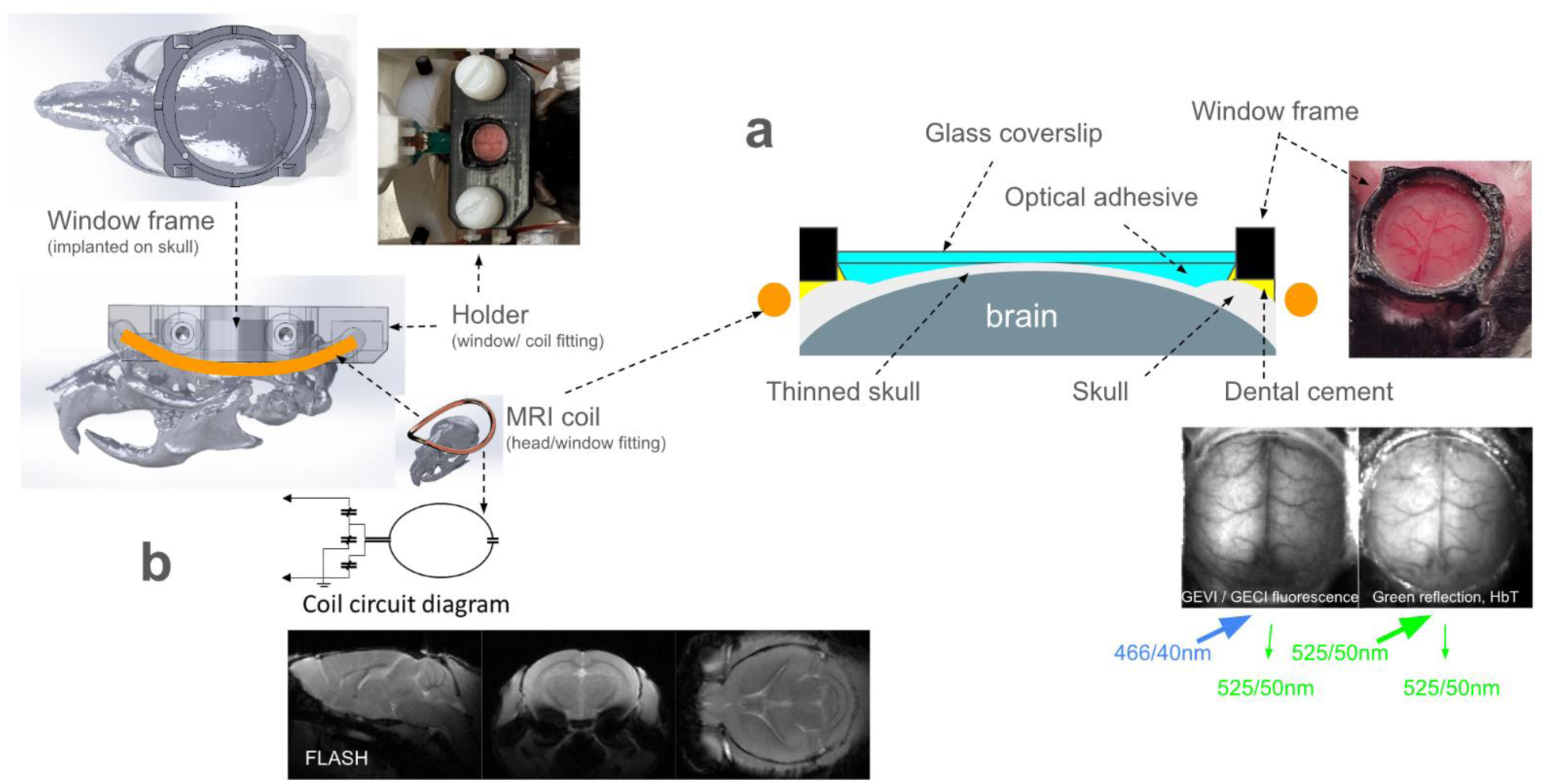
Transparent cranial window and subject-conformal RF coil. (a) Window frame bonded to skull; thinned skull replaced by optically fused glass–adhesive interface (8 mm cover glass). Interleaved fluorescence (blue excitation; green emission) and intrinsic green reflectance (HbT) are acquired. (b) Curved surface coil integrated into the head-holding assembly fits around the window frame, enabling proximity and stable positioning for whole-brain coverage.

The coil-window combination resulted in highly reproducible and stable positioning of mice. In some mice, the cranial window remained implanted for 6-12 months, but no degradation of image quality was observed, support for the feasibility of longitudinal studies. One of the mice was kept more than 15 months but image quality or FC findings are still acceptable, as shown in Figure S3.

### Simultaneous fMRI and WOI Reveal Consistent Functional Networks

To further evaluate the consistency of the cross-modality simultaneous studies at the individual level, spatial ICA was applied to fMRI or WOI, separately. All data were prepared using standard preprocessing steps, including motion correction and spatial normalization to the Allen atlas but without temporal filtering (linear detrend only), to avoid making a prediction of frequency-specific contributions to network activity. The neural circuitry of the barrel cortex and their functional connectivity (FC) in different cortical regions have been well established ^15^. As expected, both WOI and fMRI detected functional networks across hemispheres, such as bilateral barrel-field FC networks in both GECI and GEVI anmals demonstrated in Figure 3 a and b. Networks from fMRI were projected to the cortical surface using a logarithmic weighting to facilitate comparison to WOI. Our results showed similar findings comparable to previous reports across different modalities: EPI, ZTE, HbT, neural voltage or calcium. The high quality of data obtained with all three modalities is evident from the sensitivity and robustness of the results from single scans from individual mice.

**Fig. 3.**
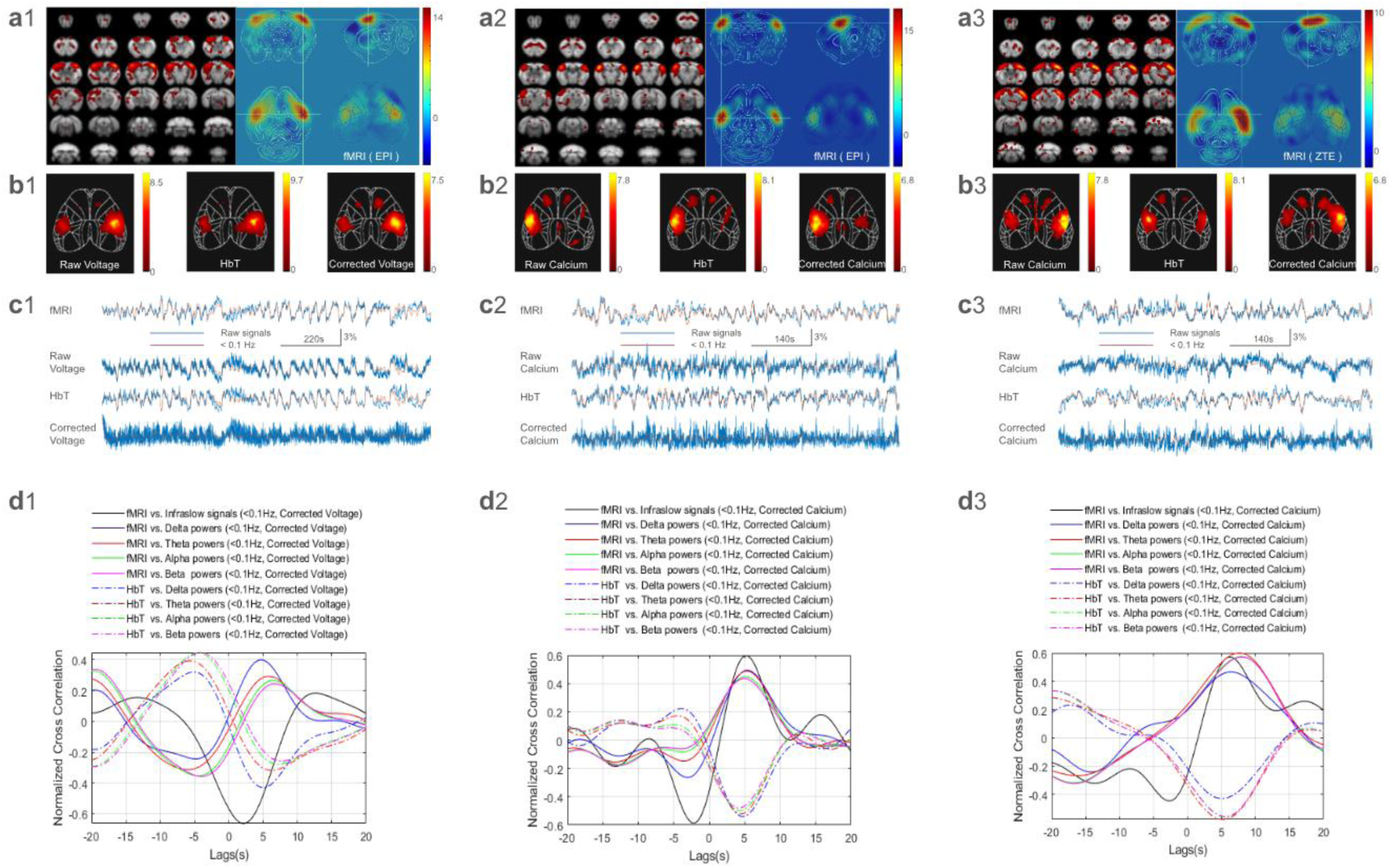
Temporal relationship between WOI and fMRI signals. Each column of panels was obtained from a single scan in an individual mouse: column 1 from a mouse with WOI of a fluorescent voltage indicator, JEDI, and fMRI with EPI; column 2 from a mouse with WOI of a fluorescent calcium indicator, GCaMP6f, and fMRI with EPI; and column 3 from a mouse with GCaMP6f and fMRI with ZTE. In rows a and b, barrel field networks detected by ICA on both fMRI (row a) and WOI (row b) are shown. As expected, networks include the barrel-field somatosensory/motor areas of the left and right hemispheres. The atlas image includes a 2D cortical projection for ease of comparison to 2D WOI cortical images (b1-3). Row c shoes the time courses (percent change, raw (blue) & lowpass filtered <0.1Hz (red)) of the detected barrel-field network from each modality. Cross correlations are compared between the hemodynamic time courses (fMRI, HbT) and fluorescence time courses were conducted with corrected fluorescent signals (d1-3).

### Temporal Analysis Dissects Hemodynamic and Neuronal Contributions to fMRI Signals

Simultaneous fMRI and WOI allowed estimation of the temporal relationship between concurrent fMRI signals and neuronal voltage/calcium signals in a given network. For example, signals from the barrel-field network shown in Figure 3a and b are plotted and compared in Figure 3c. The similarity of the raw fluorescence signal to the HbT signal is evidence of contamination from absorption related to hemodynamics, which becomes minimal after correction for hemodynamic crosstalk. To further examine the relationship between hemodynamic and neural signals, cross-correlations between the hemodynamic signals (fMRI, HbT) and the raw and corrected fluorescence signals (calcium, voltage) were conducted. Fluorescence signals were separated into frequency bands containing either infraslow signals or the infraslow envelope of various band-limited powers, Figure 3d-e, (because hemodynamics are inherently slow). The high-frequency bands (Delta to Beta) were converted to band-limited powers and filtered to the same infraslow band as fMRI (< 0.1 Hz). The cross correlations between the band-limited powers with fMRI exhibited a similar time lag as the corrected fluorescence correlation to fMRI.

As expected, a strong negative correlation was detected with approximately no time lag between fMRI and HbT (Fig. 4a), since they originate from the same hemodynamic processes but have different signal directions, as the fMRI response to neural activity is positive but the HbT absorption response is negative (i.e., more light is absorbed due to increased volume). Notably, the raw fluorescent signals, whether from GEVI or GECI, exhibited a similar time lag and signal direction to HbT (Figure 3d and 4a). After HbT regression, the corrected voltage (JEDI, Figure 3d1-e1) led changes in fMRI or HbT by a few seconds. As the JEDI voltage in neuronal activation resulted in negative signal changes ^16^, similar to hemodynamic absorption changes, we observed a positive correlation with HbT and a negative correlation with fMRI, but with consistent time lags. In contrast, neuronal calcium signals (corrected calcium) in the GCaMP exhibited positive signal changes. Therefore, we observed a positive correlation with fMRI at a time lag consistent with known hemodynamic delays, although for raw calcium fluorescence a negative correlation was observed and no lag was present. Similar results were obtained when the cross correlation was calculated on <0.1 Hz signals averaged from multiple segments of 400s long and 50% overlap for each scan, Figure S5. The similarity of the fluorescence and HbT time courses indicates that the hemodynamic contribution dominates the raw fluorescent signals. Regression of HbT minimizes this contamination and results in the expected direction and timing of the neural component relative to hemodynamics. Group analysis of GCaMP mice indicated a significant difference (Fig 4b, P=0.02, paired t-test) in neuronal activity lags between BOLD and ZTE(or HbT). The lags for ZTE and HbT exhibit similar range (Fig 4b4).

**Fig. 4.**
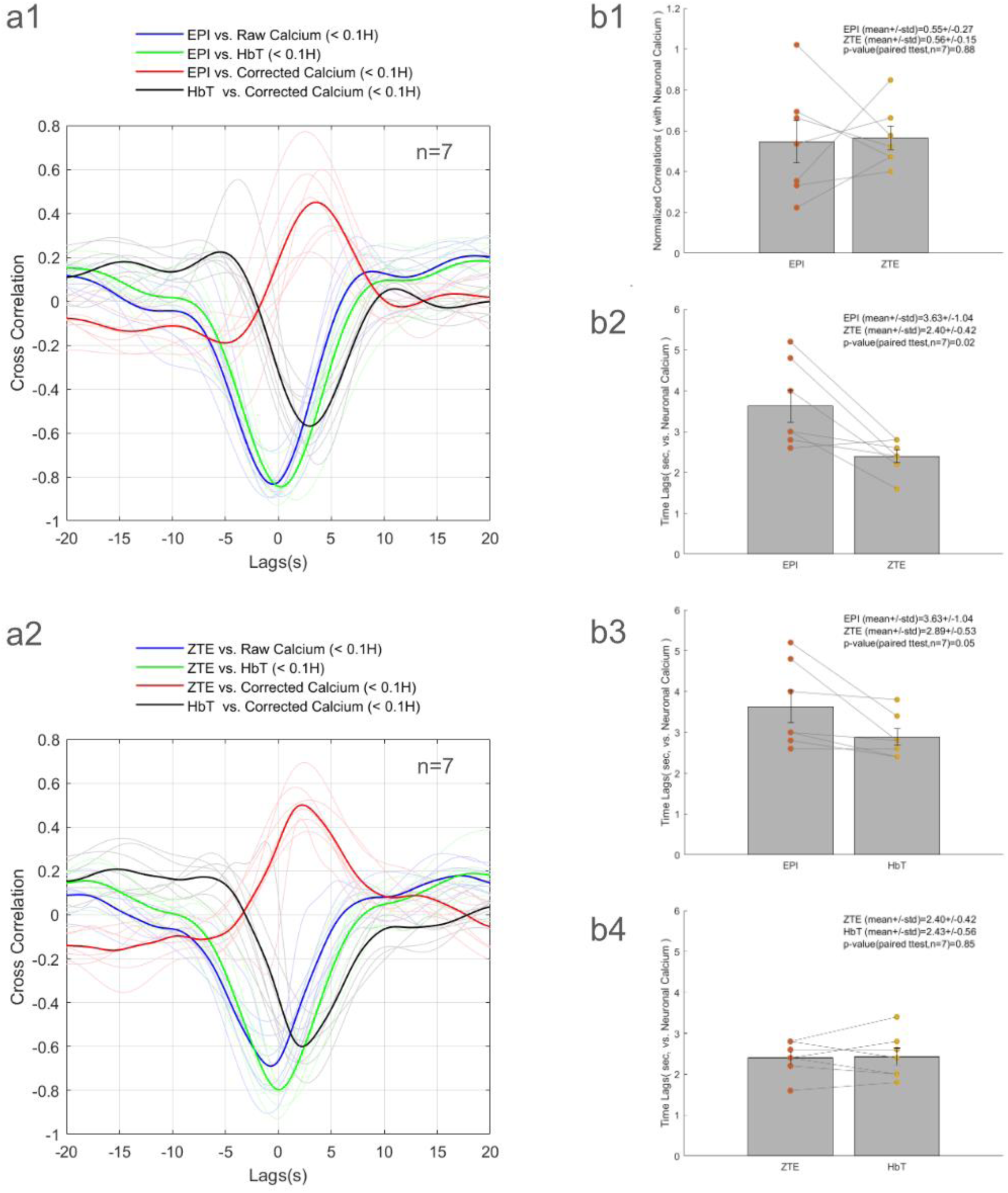
Group level comparison of temporal correlation between EPI or ZTE and concurrent infraslow neuronal calcium activities (from ICA barrel network). (a1-2) EPI or ZTE group means (solid color) and individuals (light color) of cross correlations show varied correlations and lags across raw calcium, HbT and corrected calcium. The paired ttest (n=7) indicates no significant difference for neuronal calcium correlations to EPI or ZTE (b1, p=0.88) but indicates significantly different time lags (b2, EPI>ZTE, p=0.02). The EPI time lags are also significantly longer than HbT (b3, p=0.05) but no significant difference was observed between ZTE and HbT (b4, p=0.85). Bars: mean+/-SE.

GCaMP6f has reported rise times of ~50–100 ms and decay time constants of ~200–300 ms, supporting faithful representation of neural activity up to ~4–5 Hz (Table S2). Beyond this rate, the slow decay causes individual calcium transients to merge into a single, continuous signal. JEDI voltage indicator responds on millisecond timescales, capturing up to ~50 Hz. Therefore, our frequency decomposition (infraslow <0.1 Hz; delta 1–4 Hz; theta 4–8 Hz; beta 12–25 Hz) is physiologically meaningful for calcium signals up to delta/theta bands and voltage signals across higher bands. Despite the limited information present in the calcium signal above 4 Hz, we show all frequency bands to provide context for the voltage signal. The trade-off profile between GCaMP6f and JEDI is compared in Table S2.

### Neuronal slow-to-infraslow frequencies underlie the observed FCs in fMRI

Consistent with previous reports ^2^, the infraslow frequencies (<0.1 Hz), were the dominant contributors to FC in fMRI obtained with EPI (Figure 5a1). As ZTE is a relatively new method for fMRI, FC based on ZTE is less characterized, but appears to be dominated by similar infraslow signals, Figure c1. Both fMRI mechanisms rely on neurovascular coupling to provide sensitivity to neural activity, with EPI reflecting relative oxygenation levels in hemoglobin and ZTE likely related to tissue molecular oxygen concentration, cerebral blood volume, or inflow effects (the relative contributions of which are still under investigation) ^18^. HbT exhibits infraslow frequency content, extending to ~0.5-1 Hz, consistent with the higher sensitivity and more rapid sampling of WOI measures of CBV. Corrected GCaMP6f showed intrinsic network structure from infraslow through delta bands but corrected JEDI signals exhibited structure into higher frequency bands (12-18 Hz). For both fluorescence sources, the strongest cross-modal correspondence occurred at infraslow frequencies (Figure 5a2–c2).

**Fig. 5.**
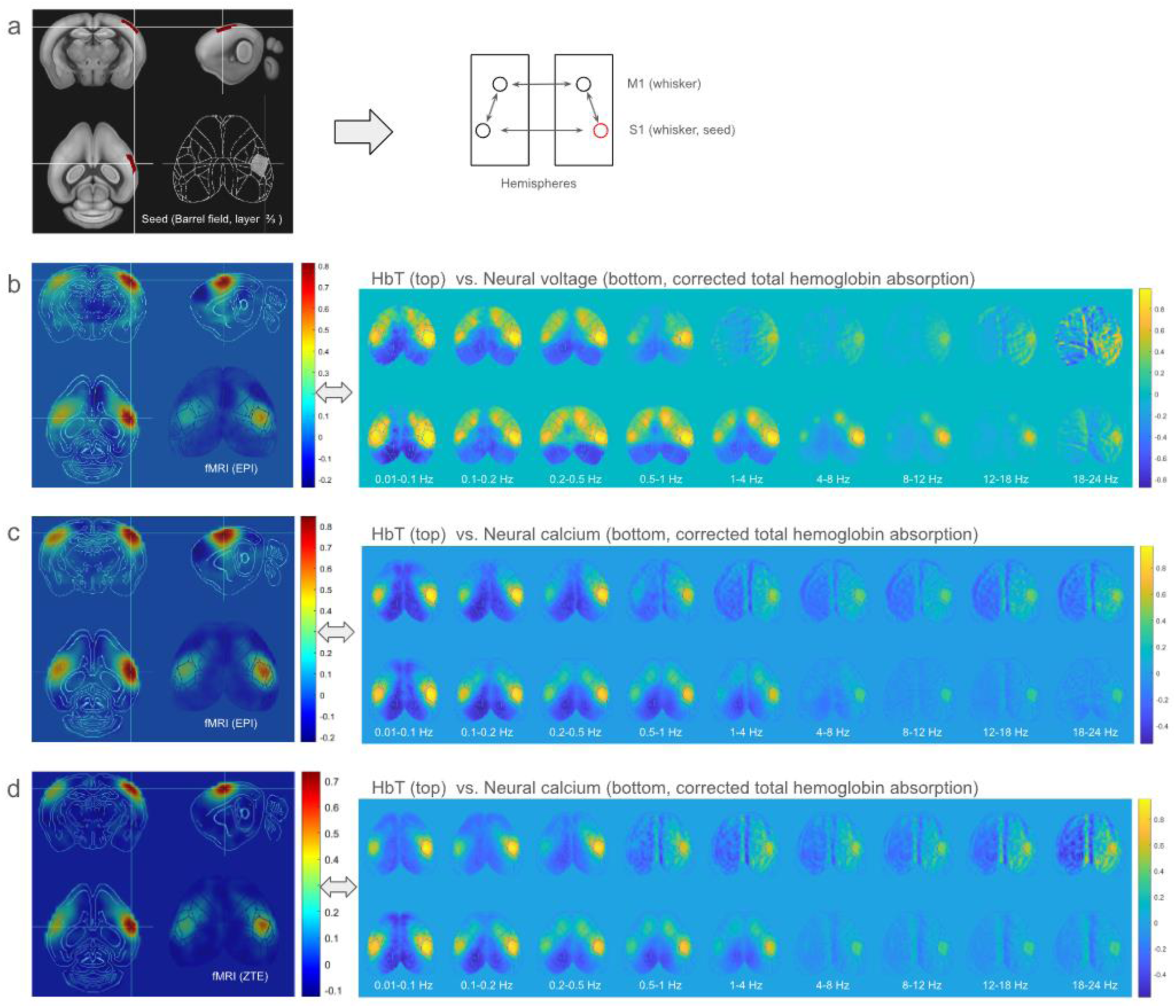
Comparison of functional connectivity (FC) between fMRI and simultaneous WOI. Group - average seed-based correlation was calculated for both WOI (voltage or calcium, 0.01–24 Hz) and fMRI (EPI or ZTE, 0.01-0.1 Hz). The WOI FCs were calculated in several frequency bands, including infraslow to slow (0.001-0.1Hz, 0.1-0.2Hz, 0.2-0.5Hz and 0.5-1Hz) and delta (1-4Hz) and theta (4-8Hz) and alpha (8-12Hz) and beta (12-18Hz and 18-24Hz). The seeds are located at 2/3 layers of the barrel field in one hemisphere (a) to detect the known network of whisker primary somatosensory of primary motor areas. (b) Simultaneous fMRI (EPI) compared with WOI (HbT and JEDI voltage, n=3), and (c) simultaneous fMRI (EPI) compared with WOI (HbT and GCaMP6f calcium, n=10) and (d) simultaneous fMRI (ZTE) compared with WOI (HbT and GCaMP6f calcium, n=7). The most consistent bilateral FC detection is in infra-slow frequency bands between WOI (neural voltage or calcium) and fMRI (EPI or ZTE). The HbT FC findings are mostly below 0.5-1Hz. Relative to GCaMP6f calcium, the JEDI voltage detected higher frequency bands FCs, i.e up to alpha-theta bands in voltage vs. up to delta bands in calcium including whisker primary motor cortex with a seed at whisker primary sensory cortex. Intrahemispheric connections are clearly present in voltage and calcium images as frequencies above the infraslow range, but they are not seen in HbT or fMRI, which better reflect the infraslow structure of neural activity.

## Discussion

The combination of WOI and fMRI in mice has the potential to significantly advance neuroscientific research and preclinical neuroimaging studies ^7,13,19–21^. A previous comprehensive review outlines the strengths and logistical hurdles of combining whole-brain fMRI with one-photon wide-field optical techniques ^22^. Recent efforts to combine wide-field optical imaging (WOI) with functional magnetic resonance imaging (fMRI) have focused on bridging the gap between cell-type-specific cortical dynamics and brain-wide hemodynamics using fiber imaging bundles to route the WOI signals from the extremely space-limited MRI scanner to a regular scientific camera outside the magnet room, avoiding custom MRI-rated electronics ^7^. While the imaging fiber bundles offer convenient routing, transmission efficiency is typically <40% and the wavelength range is limited (>500 nm). In contrast, our lens approach (>98% transmission, 400–1000 nm) maintains sensitivity comparable to benchtop optical systems while still placing the camera outside the magnet room. The high sensitivity enabled by this approach enables consistent detection of FC in individual scans (Fig. S2 and Table S2). The high transmission offsets the reduced NA in the long tube lens system, where the objective lens is set up at a distance. A shorter tube lens system could improve NA but would require custom-built MRI-compatible camera housing and is a challenge for the highly-sensitive scientific camera used for fluorescence WOI. Our method allows the full function of a regular scientific camera to be used. However, it is also possible to use MRI-compatible cameras, as demonstrated by a study of concurrent intrinsic optical imaging of absolute concentration changes in oxygenated and deoxygenated hemoglobin and simultaneous fMRI ^23^.

Neuronal cellular signals in calcium or voltage activities imaged using modern optical fluorescence technology are well established ^16,24–26^. Concurrent neuronal fluorescence imaging allows fMRI to be directly compared to the corresponding neuronal activity using the present method. Simultaneous measurement eliminates state-dependent fluctuations in neural activity/hemodynamics that can influence the detection of FC and is essential for future studies that will directly link spatiotemporal features of the neuronal cellular and fMRI signals. For both time-averaged measures (e.g., FC) and time-resolved features (e.g., coactivation patterns), simultaneous WOI and fMRI holds immense value for improved interpretation of BOLD studies in humans. Moreover, fluorescence indicators can be targeted to a wide variety of cell types, which will provide unprecedented insight into the type of activity captured in the BOLD signal. Finally, the combination of WOI and fMRI allows the rich cortical information about neural cellular activity obtained with WOI to be placed in the context of brain-wide activity that fMRI can provide.

We demonstrate a system that has sufficiently high sensitivity for both modalities that consistent results can be obtained at the individual level. The development of the integrated optical window-RF coil for our studies allows for the simultaneous acquisition of cellular voltage/calcium fluorescent signals as well as hemoglobin absorption signals across most cortical areas during whole-brain fMRI scans for a goal of comprehensive understanding of brain function across different spatiotemporal scales ^7^. The setup ensured that hemodynamic contamination of the raw fluorescence of either GECI or GEVI was minimized efficiently. Some key findings for successful WOI/fMRI integration are discussed.

### Transparent cranial window method as an essential step for high-quality WOI during fMRI

The cranial window strategy has long been used in neuroscience to gain optical access to the brain for *in-vivo* imaging. This method involves creating a transparent window in the skull through which imaging tools such as two-photon microscopy can visualize brain structures and functions. Typically, the thinned-skull cranial window is favored over craniotomy for its minimally invasive nature and ability to preserve the cerebral environment, which is crucial for longitudinal studies and for minimizing microglial activation ^27–29^. Both enable high-quality images of the pial surface ^27,30^. These windows can be implanted chronically to enable longitudinal studies in awake, behaving mice and can be tailored to specific imaging technologies or their combinations ^31^. For optimal simultaneous fMRI and WOI, the cranial window must be maximally transparent with minimal specular reflection artifacts, minimize image artifacts and interference with RF coil placement in MRI, and allow for longitudinal studies. Previous approaches have achieved some but not all these goals. For instance, a curved glass window approach has been introduced to fully replace the skull ^32^, maintaining the maximum transparency for imaging the underlying brain tissue. However, a large open skull procedure carries a risk of cortical vessel damage and bleeding, causing potential susceptibility to artifacts in fMRI. In our study, the glass surface and thinned bone were fused together by optical adhesive. The glass replacement enhanced the thinned bone strength and resulted in a super-transparent but anti-glare flat surface because both glass and optical adhesive have a similar refractive index (Refractive Index (RI), glass −1.5, optical adhesive - 1.56 in NOA81) with cranial bone: RI range from 1.555 to 1.564 at different stages of mineralization. During this procedure, air bubbles were avoided following our protocol (online). The window procedure does not add pressure to the brain but keeps the brain in a natural shape by rebuilding the bone with transparent replacements. Our thinning procedure removed all the vessels in the bone, resulting in a clear bottom layer, thereby avoiding vessel growth from the bone during chronic studies. Moreover, the reinforcement of the thinned bone procedure enhanced the durability of chronically implanted mice. Most of the mice with cranial windows (8/10) showed no significant changes in the cranial window due to growth or infarction over the course of the experiment (6-15 months). A flat surface window may be particularly useful for minimizing reflection or glare, which affect the quality of WOI. The creation of a novel cranial window is a critical step for high-quality wide-field optical imaging suitable for longitudinal studies (typically several months after cranial window implantation).

Conventional benchtop WOI studies mostly keep the skull intact. We adapted the thinned-skull method for our simultaneous fMRI studies instead due to the need to set up WOI in the space-restricted MRI scanner, where a major limitation would be the distant camera setup and resulting compromised NA. The crystal transparent optical window method reduces the light loss from the skull to improve sensitivity. A comparison of WOI in an animal with an intact skull versus our transparent rebuilt skull on thinned bone is shown in Figure S2. Both window methods were compared using benchtop WOI or simultaneous WOI/fMRI with identical animals in our pilot studies. We found similar image quality and FC detection (Figure S2) although there was a large difference in the objective distance, 4cm on-bench vs. 80cm in scanner (i.e. 20-fold NA difference when using same size objective lens). Notably, relative to the intact bone preparation, the optical window on thinned bone avoids spatial blurring due to the ‘diffusing lens’ of the intact skull.

### Unique coil combination with optical cranial window for sensitive fMRI

A subject-conformal MRI coil was specifically designed in our studies to minimize the distance between the coil and the mouse brain, and to fit closely to the region of interest, which is crucial for enhancing the signal-to-noise ratio (SNR) in MRI imaging ^33,34^. The detection sensitivity of a RF coil significantly decreases as the distance to the sample increases ^35^. In our setup, a conventional head-holding bar was used as a removable piece. Therefore, the space surrounding the optical window was clear, allowing a coil loop to fit around the window frame, close to the skin. The subject-conformal coil setup offers a direct and effective means of improving SNR because of its proximity to the mouse brain and its tailored design for specific anatomical regions. Notably, the RF Coil design was specifically engineered to handle the short-pulse requirements of ZTE (high power delivery efficiency), which standard coils might struggle with, and enhances inflow effects that contribute to the ZTE signal ^18^. The subject-conformal RF coil was positioned closely to the brain. The spatial proximity and conformal curvature is beneficial for whole-brain signal detection because it shorten the distance between the coil and subcortical areas relative to a conventional surface flat coil ^36^. In our studies, both the cranial window and the coil holder were made of 3D-printed resin and nylon material that gave no visible signals during ZTE imaging; therefore no systematic artifact from the coil or window was observed across all mice. Some materials that may have strong signals in ZTE, such as super glue and artificial tears, should be avoided.

### Efficient long tube lens solution for conventional camera imaging outside of the magnetic field

Using conventional benchtop optical imaging systems, a bulky scientific-grade camera must be set several meters away from the MRI scanner as they are typically not designed to work in a strong magnetic/RF field, ideally being placed outside the magnet room to minimize interference. As a practical solution, we demonstrate a high-quality but cost-efficient tube lens method (with no expensive image fiber bundles). With an image fiber bundle, the objective lens could set close to brain that achieves higher NA; however, the image fiber bundle typically has a limited transmission efficiency (40%) and a limited wavelength range (> 500 nm). Our tube lens solution achieves >98% transmission efficiency with a broad wavelength range (>400 nm) and is a key step toward achieving the sensitivity required to obtain consistent results from individual animals.

In addition to maximizing sensitivity, our system is designed for flexible application across a wide range of experiments. The customized 3-to-2 fiber bundle illumination method allows multiple camera channels to be employed for the simultaneous acquisition of multiple wavelengths along with the acquisition of reflectance images of intrinsic optical signals at the same wavelength. The cradle is designed for use with either anesthetized rodents on a water-circulating heating pad module or awake mice, with the heating pad module replaced by a treadmill module. The animal’s facial expression and pupil diameter are continuously monitored with a 10k-fiber endoscope (5-m long, camera outside the magnet room). For broad applications, we demonstrate simultaneous fMRI and WOI with a standard fMRI sequence (echo-planar imaging, EPI) and a more recent approach using a zero-echo time (ZTE) sequence, which has substantial advantages for deep-brain stimulation studies with less susceptibility artifacts due to electrodes and surgical damage, and for awake mice with less acoustic noise and sensitivity to head motion relative to EPI.

### Reliable representation of neuronal cellular infraslow signals in fMRI

Infraslow neuronal activity, typically defined as frequencies below 0.1 Hz or 0.5 Hz, has historically been less recognized in traditional neurophysiology, largely due to technical limitations and the predominance of faster, more easily detectable, or “classical” rhythmic EEG signals (e.g., alpha, theta, delta). However, recent research highlights that these slow direct current (DC) shifts are crucial, widespread, and potentially more representative of the underlying neural state than faster frequencies ^2,37^. Infraslow activity acts as a “carrier” for faster EEG frequencies, modulating the amplitude of higher frequency oscillations and dictating overall neuronal excitability. They are associated with behavioral performance, arousal, and the modulation of resting-state networks (RSNs). Infraslow oscillations are crucial in pathological conditions, such as epilepsy, where they are associated with the timing of interictal discharges. They are observed across the whole cortex and appear to be present in both sleeping and waking states ^38^. However, the neuronal infraslow signals can be challenging to reliably detect because they share the same frequency band with concurrent hemodynamics. Our results indicated that raw fluorescence signals were primarily dominated by hemodynamic absorption, making it necessary to minimize the modulation of hemoglobin absorption. After regressing out the hemodynamic signals, we demonstrated that fMRI time lags and corrected BOLD/ZTE signal directions become comparable to those observed in previous studies, indicating successful minimization of the hemodynamic contribution to the resting state WOI studies. Although EPI and ZTE were in similar strong correlation with concurrent neuronal activities but the time lags to the corrected neuronal calcium differ significantly (Figure 4, p=0.02). The ZTE time lags exhibited a significant short than EPI but in similar time lags with HbT that may reflect a possible contribution from CBV effects in ZTE ^39^. Future studies with higher temporal resolution and arterial spin labeling comparisons could further disentangle these contributions. With simultaneous studies, both WOI and fMRI, coordinated activity is reliably observed in the infraslow frequency bands (< 0.1 Hz). Neuronal electrophysiological infraslow activity may also play an important role in driving fMRI signals ^2,41^.

Under isoflurane anesthesia, neurons may be potentially forced into slow-wave, highly correlated, synchronous ensemble oscillations (~1 Hz) ^17^ Our studies used 1% isoflurane and observed some dominant frequency bands in FC of infraslow to delta bands with GCaMP6f and infraslow to theta bands with JEDI. These higher-frequency FCs (Figure 3 and 5) were less affected by HbT regression, indicating that they were less affected by hemodynamic absorption. We found a few seconds of time lags of fMRI to neuronal calcium activities that are consistence with previous reports under similar anesthetic conditions ^40^.

WOI and fMRI have inherently different sensitivities. WOI is mostly sensitive to the surface layers of the cortex, while fMRI is volumetric. To partially account for this difference, fMRI signals were projected across layers 1-6 using nonlinear averaging with a logarithmic decay. Nevertheless, spatial and temporal resolution are higher in HBT than in fMRI, which contributes to improved sensitivity. We expect that faster HbT FC (up to ~0.5-1 Hz) reflects improved sensitivity from the much larger number of samples enabled by the high temporal resolution, while fMRI EPI/ZTE signals are much more limited by both sampling rate and overall sensitivity.

### Future applications and biological questions

The advantages of simultaneous fMRI/WOI will enable numerous future applications for a wide range of biological questions. Functional MRI serves as a bridge between research in animal models and in human patient populations, making simultaneous fMRI/WOI an essential tool for linking the rapidly developing field of WOI transgenic mice using cellular-specific fluorescence ^42^ to human patient populations. As one example, it can often be difficult to tell whether alterations in FC in a patient population such as Alzheimers arise from changes in neural activity, hemodynamics, or both. Simultaneous WOI and fMRI can disambiguate these alterations. For fundamental neuroscience, this platform enables investigations into how specific neuronal populations (e.g., via GEVI in excitatory neurons) contribute to large-scale fMRI networks. The integrated WOI/fMRI system also allows causal testing via optogenetics of whether manipulating infraslow cortical rhythms alters whole-brain fMRI connectivity. The combination with ZTE is especially appealing for perturbation studies (e.g., deep brain stimulation, optogenetics) because it exhibits minimal artifacts ^18^.

### Limitations

The limitations that apply separately to the individual modalities remain in effect for the combined system (e.g, indicator-specific sensitivity constraints such as GEVI SNR and illumination dose, ZTE temporal SNR/undersampling trade-offs). The restriction of WOI to the cortex means that no direct subcortical neuronal readout is available for comparison with fMRI. Anesthesia may change the neurovascular response profile and certainly reduce fMRI detection sensitivity, although it remained detectable in our studies. The effects of anesthesia on neural and hemodynamic dynamics typically alter hemodynamic lags compared to awake conditions ^40^. Future work will extend to awake mouse imaging, causal perturbations (optogenetics/DBS), and targeted cell-type indicators to further resolve neuronal contributors to whole-brain fMRI.

In conclusion, the newly developed fMRI/WOI method, with a novel cranial window/RF coil design, tube lens solution, and unified data processing/registration ensures seamless integration of the two modalities and providing continuous opportunities to enhance neuroimaging. The integrated fMRI/WOI method can accommodate cellular type-specific voltage/calcium imaging, thereby paving the way to an era of precise preclinical neuroimaging. To facilitate establishing the method in the neuroimaging community, the present report includes the experimental protocol, the related 3D-printed part files, and data analysis codes.

## Methods

### Experimental animals

All animal experiments were performed in compliance with NIH guidelines and approved by the Emory University Institutional Animal Care and Use Committee. All procedures were performed on Thy1-GCaMP6f mice (Jax 024339) (n=10, five males and five females), JEDI mice ^43^ (Emx1-cre, n=3, 2 male and 1 female), and VSPF mice ^44^ (TITL-VSFPB-cre, n=1, male). The animals had access to food and water ad libitum, and were maintained under temperature-, humidity-, and light-controlled conditions.

### Cranial window and surgical procedure

Considering the balance between the maximum cortical window area and minimum RF coil size for whole-brain coverage, a cranial window frame was designed using CAD software (Solidworks, www.solidworks.com). The interior diameter of the optical window was approximately 10 mm, covering most mouse cortical regions. Files for the 3D-printed parts for both the cranial window frame and its removable holding bar are available to download along with the detailed surgical protocol (https://www.protocols.io/). Briefly, cranial surgeries were conducted in mice under 1.5-2% isoflurane anesthesia via subcutaneous injection with 2 mg/kg meloxicam SR to prevent pain and infection. The skin was removed, and the skull was exposed between the surrounding muscle attachment lines. The 3D-printed slim-profile window frame was firmly attached to the skull by using dental cement (C&B Metabond). The edges of the skin openings were closed using dental cement. The bone of the skull within the window was thinned to the bottom layer and the bone vessels were gradually and carefully removed using a dental ball bit. A UV-curing optical adhesive (NOA81, Thorlabs) and a glass coverslip (502041, WPI) were used to replace the removed bone, resulting in a super-clear and anti-glare transparent window for optical imaging of a wide field of cortical areas. The surgery lasted for approximately 2 h. Mice that underwent surgery fully recovered after 1–2 weeks and were ready for multiple months of imaging experiments under consistently good window conditions.

### Simultaneous WOI and fMRI

For simultaneous fMRI and WOI studies, it is necessary to ensure that the optical window does not interfere with the placement of the RF surface coil used to transmit and receive fMRI signals. In our system, the 3D-printed window frame fits just inside an oval, curved RF surface coil (ID, 10 mm × 15 mm) designed to fit closely around the brain to maximize sensitivity for fMRI, which is attached to the bottom of the holding bar. The lateral wings of the coil were set at approximately the center of the mouse brain for maximum whole-brain coverage parallel to the main static magnetic field (B0), ensuring that the coil RF magnetic field (B1) was perpendicular to B0 for MRI imaging. For WOI, the green GEVI/GECI fluorescence is excited by blue light (466/40 nm) and the HbT is obtained through reflectance of with green light illumination (525/50 nm). Fluorescent emission and HbT reflectance are detected at the same wavelength (525/50 nm). The holder is used to fix the head in place or release it by using two nylon screw pins from each side of the window frame. A customized curved surface coil was built into the head holder, which fits the window frame and is positioned for optimal whole-brain coverage with fMRI. The coil circuit includes matching/tuning and an optimized length of coaxial cable (~1.3-m) connecting to the scanner preamplifier. The MRI image quality and brain coverage with the coil setup is demonstrated in anatomical imaging with FLASH in Figure 2. The optical imaging tube lens system was designed to fit inside the magnet and was connected to a customized simultaneous multi-camera system set outside the magnet room ^13^. A multichannel fiber bundle running to both sides of the cranial window provided illumination at 45-degree angles from the back bilaterally. The blue LED light was filtered (Semrock, FF01-466/40) for fluorescence excitation. The emitted green-fluorescent signals were filtered (Semrock, FF03-525/50) before entering a high-speed CMOS camera (KURO 1200 B). The camera channel was also used to capture reflected intrinsic optical signals (IOS) at the same wavelength, alternating at 50 Hz over direct green light illumination with alternating 50 Hz for fluorescent green images. The image resolution was 100 × 100 pixels on a 16-bit grey scale image. Simultaneous fMRI scans were conducted using a Bruker BioSpin 9.4T scanner with an AVANCE NEO console and Paravision360 v3.5. A slim and curved surface coil with compact match/tuning circuits was customized to fit the implanted window piece, allowing close positioning to the mouse head for sensitivity. FOV saturation pulses were applied to the eye sides of the head to minimize non-brain signals. Both EPI and ZTE were set to the same spatial resolution of ~ 400-µm isotropic voxels with whole-brain coverage and a temporal sampling rate of 1s per brain volume scan. Single-shot gradient echo EPI: TR=1000ms/TE=12ms for 16 continuous axial slices. ZTE: TR, 0.673 ms; flip angle, 3°; bandwidth, 128 kHz; matrix size, 54 × 54 × 54; field of view, 22 × 22 × 22 mm^3^. The 20 min scan sessions alternated between ZTE and EPI for resting-state scans under 1% isoflurane. 2D-TOF angiography was performed to image the cortical vessel structure for registration with optical images. ECG, respiration pillow signals, and rectal temperature signals were sampled every 10ms and recorded during each fMRI session. The pupil dynamic and facial behavior were monitored by an imaging-fiber camera at 12.5 Hz as reference to control animals in a stable physiological condition.

### Data analysis

Both fMRI and WOI data were routinely preprocessed and registered in the Allen atlas using a strategy based on tissue-boundary landmarks ^45^. The preprocessing steps included (fMRI) slice timing, motion correction, top-up distortion correction, 0.5 mm-FWHM 3D smoothing, and atlas registration; (WOI) down-sampling to 2-fold of voxel size, smoothing with 0.15 mm FWHM and atlas registration. Whole-brain angiography images were obtained using MRI 2D-TOF and registered to the Allen atlas in 3D space. Cortical slices of spatially normalized vessel images were averaged and projected onto a 2D cortical Allen atlas ^46^. An individual vessel-based 2D cortical atlas was used to register the WOI images to the Allen cortical atlas (Figure S6). For comparison, in addition to 3D slices, fMRI maps were also shown in a similar way to the 2D cortical view of the camera WOI by weighted cortical layers projection^47^. We used both ROI-based and date-driven methods in our analysis with fMRI or WOI seperately. In our ICA studies (GIFT, https://trendscenter.org/software/gift/) all data were prepared using minimum preprocessing steps without predicting the frequency contribution to network detection, including motion correction and spatial normalization to the Allen atlas without temporal filtering after a linear detrend. The time courses of the barrel field networks were used for cross-correlation analysis between fMRI and WOI. For the seed-based analysis, all data were filtered in various frequency bands to evaluate the frequency contribution. The green reflection light was used to image the total hemoglobin absorption of CBV signals and was regressed from the fluorescence signal to minimize the influence of hemoglobin absorption and purify neuronal signals. To evaluate frequency contribution on FC, ROI-based analysis conducted with varied frequency bands across modalities.

Group comparisons used two-tailed paired t-tests unless stated; normality was checked (Shapiro-Wilk). Reported values are mean ± SE unless noted; No data was excluded unless pre-registered motion thresholds (>0.1 mm) were exceeded.

## Supporting information

Supplementary protocol

## Data and code availability

All data supporting the findings, key analysis codes, and 3D-print files are publicly available at Zenodo with the identifier [DOI will be added upon reviewer-access/acceptance. The cranial window surgery protocol is available (https://www.protocols.io/).

## Acknowledgements

Supported by NIH R01-NS078095, R01-MH111416, and R01-EB029857.

## References

1. Pan, W.-J. et al. Broadband Local Field Potentials Correlate with Spontaneous Fluctuations in Functional Magnetic Resonance Imaging Signals in the Rat Somatosensory Cortex Under Isoflurane Anesthesia. Brain Connectivity 1, 119–131 (2011).

2. Pan, W.-J. J. W.-J., Thompson, G. J. G. J., Magnuson, M. E. M. E., Jaeger, D. & Keilholz, S. Infraslow LFP correlates to resting-state fMRI BOLD signals. NeuroImage 74, 288–297 (2013).

3. Lu, H. et al. Synchronized delta oscillations correlate with the resting-state functional MRI signal. Proc Natl Acad Sci U S A 104, 18265–18269 (2007).

4. Brinker, G. et al. Simultaneous recording of evoked potentials and T2*-weighted MR images during somatosensory stimulation of rat. Magn Reson Med 41, 469–473 (1999).

5. Logothetis, N. K., Pauls, J., Augath, M., Trinath, T. & Oeltermann, A. Neurophysiological investigation of the basis of the fMRI signal. Nature 412, 150–157 (2001).

6. Shmuel, A. & Leopold, D. A. Neuronal correlates of spontaneous fluctuations in fMRI signals in monkey visual cortex: Implications for functional connectivity at rest. Hum Brain Mapp 29, 751–761 (2008).

7. Lake, E. M. et al. Simultaneous cortex-wide fluorescence Ca2+ imaging and whole-brain fMRI. Nat Methods 17, 1262–1271 (2020).

8. Gozzi, A. & Zerbi, V. Modeling Brain Dysconnectivity in Rodents. Biological Psychiatry 93, 419–429 (2023).

9. Aladjalova, N. A. Infra-Slow Rhythmic Oscillations of The Steady Potential of the Cerebral Cortex. Nature 179, 957–959 (1957).

10. Imamura, A. et al. Zero-echo time imaging achieves whole brain activity mapping without ventral signal loss in mice. NeuroImage 307, 121024 (2025).

11. Ljungberg, E. et al. Silent zero TE MR neuroimaging: Current state-of-the-art and future directions. Progress in Nuclear Magnetic Resonance Spectroscopy 123, 73–93 (2021).

12. MacKinnon, M. J. et al. fMRI with a Zero Echo Time (ZTE) Pulse Sequence. Program # 2910.

13. Pan, W.-J. et al. (ISMRM 2022) Optimization of wide-field optical imaging method towards fMRI integration in mice. https://archive.ismrm.org/2022/3331.html (2022).

14. Pan, W.-J., Zhou, L., Clavijo, G. P., Sharghi, V. & Keilholz, S. (ISMRM 2021) Optimized Single-loop Coil with 3D-shaped Design for Simultaneous fMRI and Optical Imaging in Rodent. https://archive.ismrm.org/2021/2511.html (2021).

15. Gellért, L., Luhmann, H. J. & Kilb, W. Axonal connections between S1 barrel, M1, and S2 cortex in the newborn mouse. Front. Neuroanat. 17, (2023).

16. Lu, X. et al. Detecting rapid pan-cortical voltage dynamics in vivo with a brighter and faster voltage indicator. 2022.08.29.505018 Preprint at 10.1101/2022.08.29.505018 (2022).

17. Xu, R., Chen, R., Zhang, X., Zhang, X. & Zhang, J. Neuronal activity in the cortex of mice under anesthesia depend on the type of anesthetic drug. BMC Anesthesiol 25, 452 (2025).

18. Mangia, S., Michaeli, S. & Gröhn, O. Outlook on zero/ultrashort echo time techniques in functional MRI. Magn Reson Med 95, 714–723 (2026).

19. Kennerley, A. J., Mayhew, J. E., Boorman, L., Zheng, Y. & Berwick, J. Is optical imaging spectroscopy a viable measurement technique for the investigation of the negative BOLD phenomenon? A concurrent optical imaging spectroscopy and fMRI study at high field (7 T). Neuroimage 61, 10–20 (2012).

20. Kennerley, A. J. et al. Refinement of Optical Imaging Spectroscopy Algorithms using concurrent BOLD and CBV fMRI. Neuroimage 47, 1608–1619 (2009).

21. Kim, S. et al. Whole-brain mapping of effective connectivity by fMRI with cortex-wide patterned optogenetics. Neuron 111, 1732–1747.e6 (2023).

22. Lake, E. M. R. & Higley, M. J. Building bridges: simultaneous multimodal neuroimaging approaches for exploring the organization of brain networks. Neurophotonics 9, 032202 (2022).

23. Bernard, R., Valverde Salzmann, M., Scheffler, K. & Pohmann, R. Concurrent intrinsic optical imaging and fMRI at ultra-high field using magnetic field proof optical components. NMR in Biomedicine 36, e4909 (2023).

24. Russell, J. T. Imaging calcium signals in vivo: a powerful tool in physiology and pharmacology. Br J Pharmacol 163, 1605–1625 (2011).

25. Knöpfel, T. & Song, C. Optical voltage imaging in neurons: moving from technology development to practical tool. Nature Reviews Neuroscience 20, 719–727 (2019).

26. Lu, X. et al. Widefield imaging of rapid pan-cortical voltage dynamics with an indicator evolved for one-photon microscopy. Nat Commun 14, 6423 (2023).

27. Li, Y., Baran, U. & Wang, R. K. Application of Thinned-Skull Cranial Window to Mouse Cerebral Blood Flow Imaging Using Optical Microangiography. PLOS ONE 9, e113658 (2014).

28. Marker, D. F., Tremblay, M.-E., Lu, S.-M., Majewska, A. K. & Gelbard, H. A. A Thin-skull Window Technique for Chronic Two-photon In vivo Imaging of Murine Microglia in Models of Neuroinflammation. J Vis Exp 2059 (2010) doi:10.3791/2059.

29. Yang, G., Pan, F., Parkhurst, C. N., Grutzendler, J. & Gan, W.-B. Thinned-skull cranial window technique for long-term imaging of the cortex in live mice. Nat Protoc 5, 201–208 (2010).

30. Isshiki, M. & Okabe, S. Evaluation of cranial window types for in vivo two-photon imaging of brain microstructures. Microscopy 63, 53–63 (2014).

31. Kılıç, K. et al. Chronic Cranial Windows for Long Term Multimodal Neurovascular Imaging in Mice. Front. Physiol. 11, (2021).

32. Kim, T. H. et al. Long-Term Optical Access to an Estimated One Million Neurons in the Live Mouse Cortex. Cell Reports 17, 3385–3394 (2016).

33. Vanduffel, H. et al. Additive Manufacturing of Subject-Conformal Receive Coils for Magnetic Resonance Imaging. Advanced Materials Technologies 7, 2200647 (2022).

34. Hadley, R., Parker, D. & Minalga, E. Improving signal-to-noise ratio in transcranial magnetic resonance guided focused ultrasound. J Ther Ultrasound 3, O36 (2015).

35. Kwok, W. E. Basic Principles of and Practical Guide to Clinical MRI Radiofrequency Coils. RadioGraphics 42, 898–918 (2022).

36. (ISMRM 2021) Optimized Single-loop Coil with 3D-shaped Design for Simultaneous fMRI and Optical Imaging in Rodent. https://archive.ismrm.org/2021/2511.html.

37. Vanhatalo, S. et al. Infraslow oscillations modulate excitability and interictal epileptic activity in the human cortex during sleep. Proceedings of the National Academy of Sciences 101, 5053–5057 (2004).

38. Mitra, A. et al. Spontaneous Infra-slow Brain Activity Has Unique Spatiotemporal Dynamics and Laminar Structure. Neuron 98, 297–305.e6 (2018).

39. Mangia, S., Michaeli, S. & Gröhn, O. Outlook on zero/ultrashort echo time techniques in functional MRI. Magnetic Resonance in Medicine 95, 714–723 (2026).

40. Ma, Y. et al. Resting-state hemodynamics are spatiotemporally coupled to synchronized and symmetric neural activity in excitatory neurons. Proceedings of the National Academy of Sciences 113, E8463–E8471 (2016).

41. Grooms, J. K. et al. Infraslow Electroencephalographic and Dynamic Resting State Network Activity. Brain Connectivity 7, 265–280 (2017).

42. Arias, A., Manubens-Gil, L. & Dierssen, M. Fluorescent transgenic mouse models for whole-brain imaging in health and disease. Front Mol Neurosci 15, 958222 (2022).

43. Lu, X. et al. Widefield imaging of rapid pan-cortical voltage dynamics with an indicator evolved for one-photon microscopy. Nat Commun 14, 6423 (2023).

44. Meyer-Baese, L. et al. Cortical Networks Relating to Arousal Are Differentially Coupled to Neural Activity and Hemodynamics. J. Neurosci. 44, (2024).

45. Pan, W.-J., Anumba, N., Xu, N., Meyer-Baese, L. & Keilholz, S. (ISMRM2023) Manual registration and customized template for rodent fMRI data spatial normalization. https://www.ismrm.org/23/program-files/D-19.htm.

46. Pan, W.-J., Daley, L., Watters, H., Meyer-Baese, L. & Keilholz, S. (ISMRM 2024) Investigating concurrent neuronal activities and hemodynamic signals with simultaneous fMRI and wild-field cortical optical imaging in mice. ISMRM2024 https://submissions.mirasmart.com/ISMRM2024/Itinerary/PresentationDetail.aspx?evdid=3608.

47. Pan, W.-J., Daley, L., Watters, H., Meyer-Baese, L. & Keilholz, S. Tracing neuronal synchronized slow oscillations with simultaneous fMRI and optical imaging in mice. in (Seoul, 2024).

