## Supplementary protocol for "An integrated platform for simultaneous wide-field voltage/calcium imaging and fMRI (EPI & ZTE) reveals neuronal infraslow dynamics underlying functional connectivity"

**Protocol** (online, https://www.protocols.io/ )

Abstract

For successful simultaneous wide-field optical imaging (WOI) during fMRI in mice, a specific cranial window design is required for high-quality imaging of both WOI and fMRI. For this purpose, we developed a protocol for building a cranial window that integrates a custom head holder and subject-conformal RF surface coil. The transparent cranial window method included nine sections and step-by-step details. Files for all 3D printed parts are shared along with the methods for quick adoption in the neuroimaging community.

Section 1.

3d-printed parts preparation. Designed in SolidWorks (part files attached) and 3D printed with resin.

1. Cranial window frame (resin, wframe.SLDPRT).

2. Cranial window positioning tool (resin, ptool.SLDPRT).

3. Holding bar (resin, hbar.SLDPRT, 2-56 thread late) and screw pins (45-degree tip conical shape of 2-56 nylon screw).

Section 2.

Sterile Preparation

1. Autoclave surgical instruments, drill bits, headpieces, opened sterile paper tissue, non-sterile clear wrappers, mixing stickers, ceramic mixer (for dental cement) in an autoclave pack with drape-wrapped and sealed autoclave tape indicator. A steam indicator strip or piece of autoclave steam tape should also be placed inside the package.

Section 3. Surgery Area Preparation

1. Turn on the heating pad to reach and remain at 39 °C by the feedback from on-pad sensor before setting the animal on it. Be sure the temporal sensor is attached to the heating pad properly to monitor the pad temperature as setting.

2. Check the isoflurane level in the vaporizer and fill the vaporizer with isoflurane, if needed.

3. Weigh the charcoal canister and record the current date and weight on the side of the canister (discard and replace if the canister gained 50 g from the initial weight).

4. Turn on oxygen supply.

5. Wipe all surgical surfaces with DAR-provided disinfectants.

6. Place the lab paper tissue covering the heating pad.

7. Place an autoclaved pack, eye ointment, hair clipper, hair depilatory lotion, cotton-tipped applicators, double-sided tape, etc.

8. Place a clean cage halfway on the heating pad for animal recovery.

9. Start filling in the Surgery Log and IACUC Surgery Card.

Section 4. Surgeon Preparation

1. Wash hands.

2. PPE: lab coat (completely buttoned) or disposable gowns.

3. Surgical-type mask.

4. Exam gloves.

5. Hair cover.

Section 5. Drug Preparation

1. Meloxicam SR 2–4 mg/kg, 21–22-gauge needle with a 1cc syringe.

Section 6. Animal Preparation

1. Weigh the animal and record the weight in the surgery log and surgery card.

2. Adjust oxygen flow to 1 LPM and switch flow to an induction chamber.

3. Place the animal in the induction chamber, turn on the vaporizer dial to 3-3.5% for mice, and set a timer for 3 min.

4. Monitor the animal for respiration rate (down to approximately 1 Hz), typically in 3 min.

5. Turn off isoflurane and flush chamber with oxygen 3sec by depressing the oxygen flush valve.

6. Switch anesthesia tubing to direct flow to the nose cone on the surgery table, then turn isoflurane back on to 1.5%.

7. Move animal to nosecone on the surgery table. Monitor the breathing rate visually periodically and ensure that it is above 1 breath/s during animal anesthetization.

8. Secure animals with tooth bar in the nosecone before hair removal.

9. Adjust the head height via the tooth-bar holder to ensure that breathing is not restricted.

10. Apply eye lubricant to both eyes using a cotton-tipped applicator.

11. Subcutaneous injection of prepared meloxicam SR (2 mg/kg), pinching the skin at the injection site while withdrawing the needle, and then holding the skin pinched for 5-10 seconds.

12. Remove using a clipper, ensuring that no hair is included in the closure of the incision.

13. Remove any loose hairs from the animal and around the surgical area using cotton ball and tape.

14. Apply depilatory hair lotion on the surgery area for 20-30 sec for the mouse. Remove quickly with cotton and wash any excess from the skin with water to avoid chemical burns.

15. Set mouse ear bars in place, adjust head height to avoid any restriction of breathing.

16. Skin antiseptics: apply alcohol/iodine pads three times alternatively, ending with iodine.

17. Change into a new pair of exam gloves.

18. Open the surgical pack for drapes and scissors.

19. Cut a hole of surgery area size for an autoclaved clear drape with autoclaved scissors, covering the entire animal. Be sure that the fenestration hold does not expose hair. Double-sided tape can be used on the surrounding metal surface of a stereotactic instrument to hold the drape in place.

Section 7. Surgical procedure for cranial window frame implantation

1. Arrange the surgical instruments on sterile paper tissue to maximize sterility maintenance, handles toward the surgeon, and place all instrument tips above an imaginary line to remain sterile. Be sure using the sterile tip technique; only touch the handles of the instruments with examination gloves.

2. Before making the first incision, gently pinch the foot with the instrument through the top of a clear drape. If animals show any response, increase isoflurane by 0.5%, wait for 1 min, and test foot pinch again.

3. Begin surgery once a lack of pain response is confirmed. Monitor respiration at least 60 breaths/min.

4. Cut off the skin overhead, slightly larger than the prepared window headpiece.

5. Clean tissue over the skull with sterile cotton swabs and sterile needles.

6. Keep the skull surface clean dry.

7. Place the autoclaved dental cement mixer on an ice-bed container.

8. Mix the C&B Metabond on the chilled mixing dish. Four drops of Metabond liquid base and one drop Metabond catalyst mixed with a Metabond power part.

9. Quickly apply a thin layer of the mixed Metabond on the skull and the opened skin edges.

10. Remove the surgical drape and adjust the ear bar for level head positioning if needed.

11. Attach the prepared window piece to a calibration tool (optional) and place the bottom-curved headpiece on the skull closely, ensure correct positioning based on skull anatomical marks, middle suture line, Bregma/Lambda points, holding on by the calibration tool. Apply Metabond further between the window frame piece and skull carefully, ensuring no gaps or air bubbles.

Section 8. Construct a transparent cranial window.

1. Thin-bone procedure in the implanted window: Under the surgical scope, the layers of bone were removed gradually, but slowly and carefully, by ball milling with a dental drill. All vessels in the bone, which are mostly in the middle layers, are removed for optimal optical properties. Ensure no damage to the deep bone layer or the dura. Small spot bleeding from bones may be common but can be easily stopped by milling. An air puffer may be used to remove the milled bone powders for a clear view periodically during the procedure.

2. Apply a thin layer of Metabond on the thinned bone for enhancement.

3. Place an 8 mm-diameter glass coverslip over the thinned bone.

4. Tilt down the mouse head at approximately 10 °and position the glass window in the center of the frame.

5. Drop optical adhesive from lower side of the glass window. The optical adhesive will fill up between the bone and glass coverslip without air bubbles trapped in.

6. Tilt back mouse head in level. Check and ensure no bubbles and adjust glass coverslip positioning in center in need.

7. UV cure over a pin-hole plate. Moving the pin-hole plate between the UV beam and the cranial window for gradually curing up all window areas without generating too much heat or unbalanced heating induced volume changes.

8. If there is any adhesive on glass, use a 25G needle to clean up the surface under a surgical scope.

Section 9. Concluding surgery

1. Carefully check for any gaps between skin and headpiece, be sure no open skin and no blood.

2. Turn off isoflurane.

3. Place the animal in the prepared clean recovery cage that is halfway on the heating pad.

4. Monitor the animal every 15 minutes until you walk around the cage, document the animal's condition on the surgery log.

5. Complete surgery log and IACUC surgery card.

6. Set the surgery card with a cage card.

7. Return the animal to the new home cage with dampened mouse chow in a small dish and return to the animal room.

8. Clear the surgery area and surgery instruments using DAR antiseptic spray.

9. Subcutaneous injection of meloxicam SR (2mg/kg) per day for continuous 3 days and recording animal condition.

10. After 1-2 weeks of recovery, the animal will be ready for functional imaging studies.
