## Supplementary figures and tables for "An integrated platform for simultaneous wide-field voltage/calcium imaging and fMRI (EPI & ZTE) reveals neuronal infraslow dynamics underlying functional connectivity"

### Supplementary materials

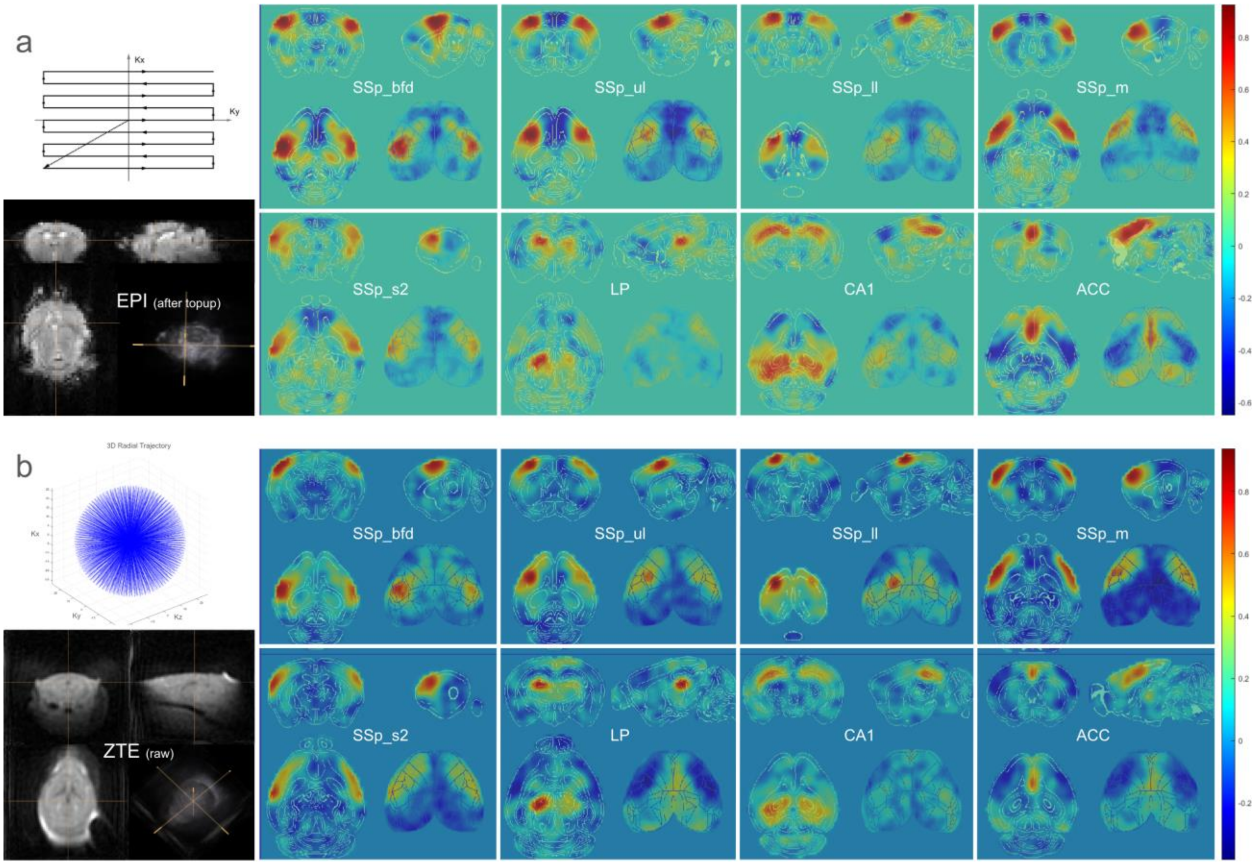


**Fig. S1** Example of FC for EPI and ZTE within a representative subject. Seed-based correlation analysis was conducted from preprocessed data of EPI or ZTE scans in mice anesthetized with 1% isoflurane. Raw intensity images are shown and compared for echo plane imaging (EPI) and radial trajectory zero echo time imaging (ZTE). Pearson correlation values are shown; the regions of seed locations in Allen coordinates are indicated in the maps respectively. Seeds were selected from various cortical and subcortical regions of the left hemisphere for EPI (a) or ZTE (b), including barrel-field primary somatosensory area (SSp_bfd), upper-limb primary somatosensory area (SSp_ul), lower-limb primary somatosensory area (SSp_ll), mouth primary somatosensory area (SSp_m), secondary somatosensory area (SSp_s2), lateral posterior nucleus of the thalamus (LP), hippocampus cornu ammonis 1 (CA1), anterior cingulate cortex (ACC). Comparable networks are detected with both fMRI sequences.

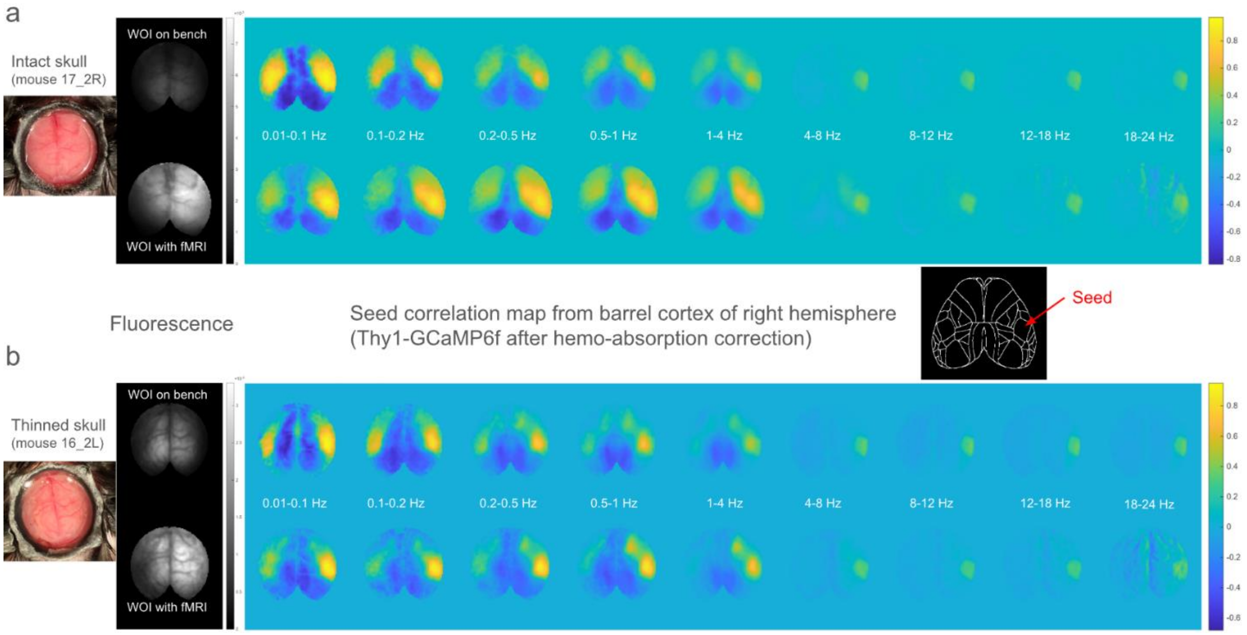


**Fig. S2** Comparison between benchtop WOI and WOI/fMRI, and between intact skull and thinned skull preparations. The pilot studies were conducted with mice from the same cage (matched on age and sex). (a) The intact skull window was prepared without thin-bone procedure but including other steps as well as thinned skull window (b), i.e. optical adhesive fusion with glass top. The thin-bone procedure removed most tissue and vessels in bone but kept the lower layer intact and shows the cortical vessels more clearly than in the intact skull, b vs. a. The intact skull may serve as a diffusing filter that blurs FC maps and reduces spatial accuracy. Comparing benchtop WOI to fMRI/WOI, both image quality and FC findings are similar despite the setup limitations with fMRI (distant objective, 80cm with fMRI vs. 4cm on bench) without SNR reduction.

| | **SNR**<br>**WOI on bench** | **SNR**<br>**WOI with fMRI** |
|---|---|---|
| Skull intact<br>(mouse 17_2R) | 22.8+/-1e-05 | 35.3+/-3e-05 |
| Thinned skull<br>(mouse 16_2L) | 26.2+/-2e-05 | 31.6+/-2e-05 |

* Note: SNR calculation: average brain SNR of repeated scans along time points, i.e. a = image at one time point, b = image at next time point, c = b-a, SNR_ = mean(a(brain)). /Std(c(brain))/sqart(2), SNR = mean (all time points of SNR_) +/- SE

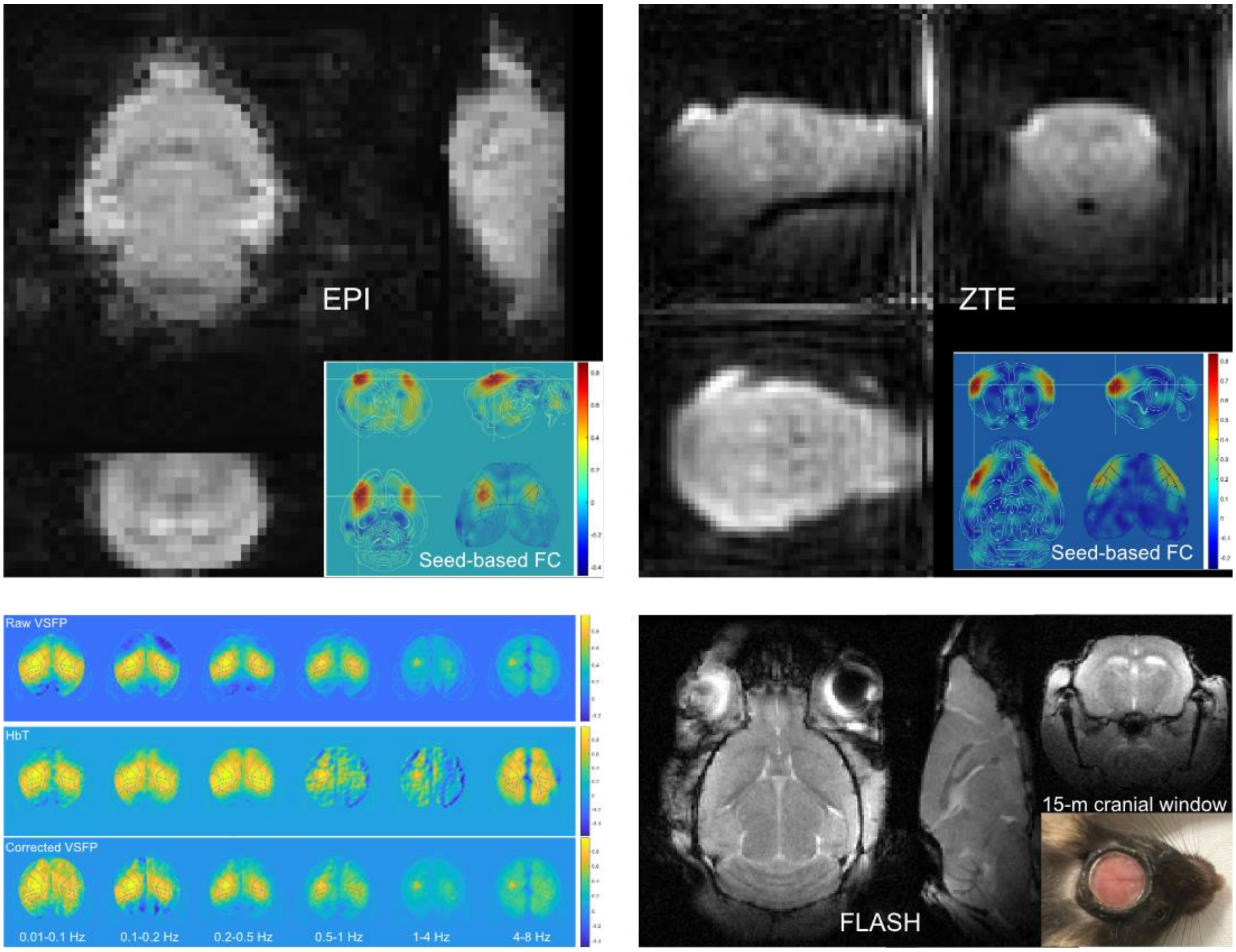


**Fig. S3** Example of long-lasting cranial window. Both fMRI (EPI/ZTE) and WOI (HbT/VSFP) appeared acceptable for a representative mouse (79R) at 15 months after cranial window implantation.

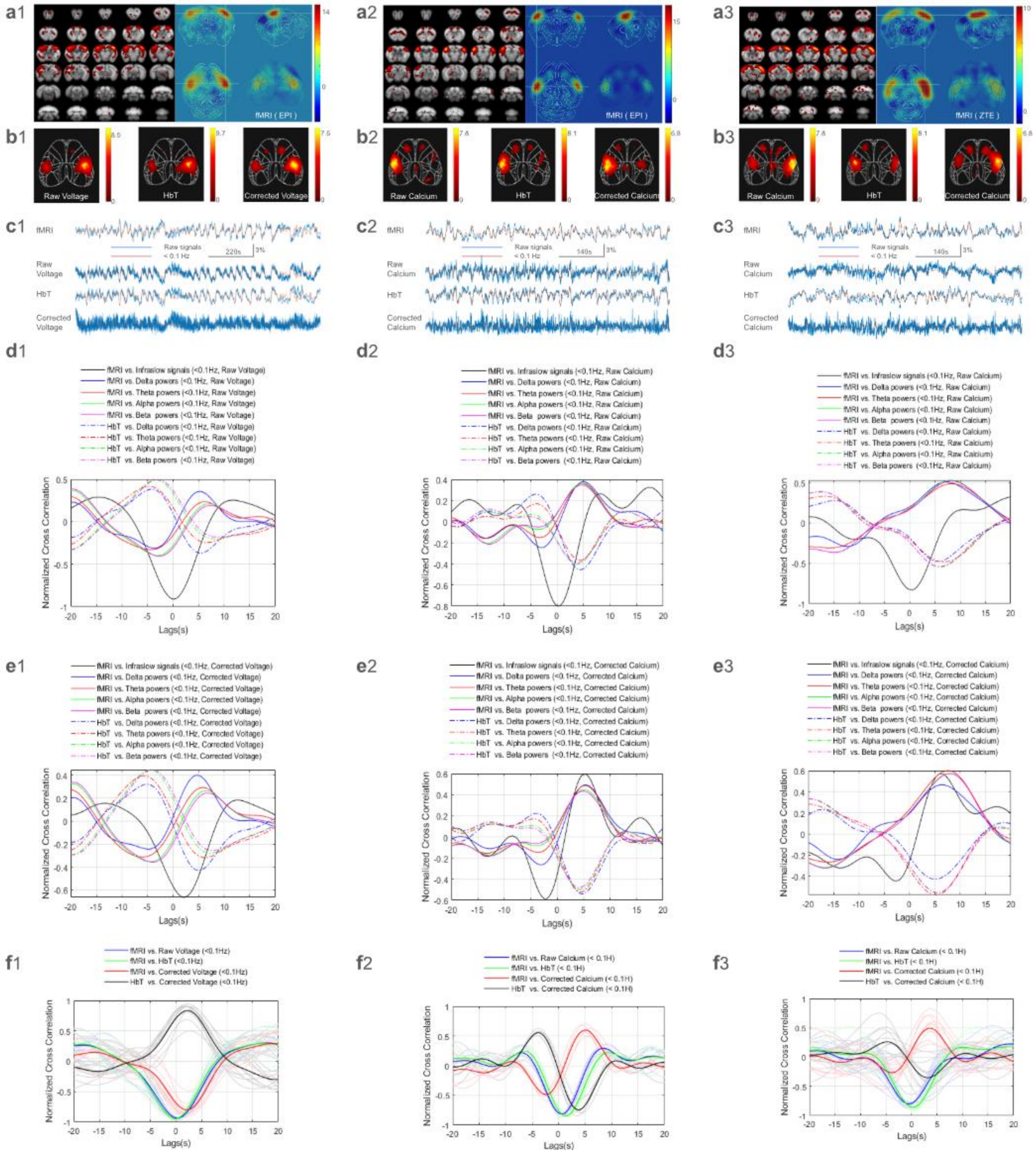


**Fig. S4**. Functional connectivity (FC) for simultaneous fMRI (EPI/ZTE) and WOI (GEVI/GECI) and the temporal relationship between WOI and fMRI signals. Each column of panels was obtained from a single scan in an individual mouse: column 1 from a mouse with WOI of a fluorescent voltage indicator, JEDI, and fMRI with EPI; column 2 from a mouse with WOI of a fluorescent calcium indicator, GCaMP6f, and fMRI with EPI; and column 3 from a mouse with GCaMP6f and fMRI with ZTE. In rows a and b, barrel field networks detected by ICA on both fMRI (row a) and WOI (row b) are shown. As expected, networks include the barrel-field somatosensory/motor areas of the left and right hemispheres. The fMRI cortical results are shown first on slices of the 3D anatomical image and then registered to the volumetric Allen brain atlas. The atlas image includes a 2D cortical projection for ease of comparison to 2D WOI cortical images (b1-3). For WOI images, the raw fluorescence results, the HbT results, and the corrected fluorescence results are shown. Row c shoes the time courses (percent change, raw (blue) & lowpass filtered <0.1Hz (red)) of the detected barrel-field network from each modality. The similarity of the raw fluorescence signal and the HbT signal is evidence of hemodynamic contamination of the raw fluorescence, which is minimized in the corrected fluorescence signal. To further demonstrate the presence and removal of hemodynamic confounds, cross-correlation between the hemodynamic time courses (fMRI, HbT) and fluorescence time courses were conducted with raw fluorescent signals (d1-3) or corrected fluorescent signals (e1-3). To allow possible variation based on the frequency content of the fluorescence signals, they were first filtered into frequency bands ranging from infraslow to beta. For the infraslow band (<0.1 Hz), the raw fluorescence signals were dominated by hemoglobin absorption (HbT), showing high similarity, i.e. strong correlation with fMRI without time lags. After HbT regression, the corrected fluorescence exhibited a few seconds of time lags, comparable to expected hemodynamic delays. Notably, GEVI and GECI have different signal changes during activity (voltage decreases but calcium increases during activity). Consistently it was observed that the corrected fluorescence in GECI changed direction from the raw GECI (d2-3 vs. e2-3) in addition to exhibiting a lag, while both the raw and corrected GEVI have same negative direction as HbT and only the lag changed after HbT regression (d1 vs. e1). The high-frequency bands (Delta to Beta) were converted to band-limited powers and filtered to the same infraslow band as fMRI (< 0.1 Hz). The cross correlations between the bandlimited powers with fMRI exhibited a similar time lag as the corrected fluorescence correlation to fMRI. For the higher frequencies, Delta to Beta, there were no differences of fMRI correlations with the raw or corrected fluorescence (GEVI or GECI with EPI or ZTE), consistent with the low frequency content of the hemodynamic contamination. Segment-based cross correlations compared between fMRI and infraslow band of GEVI or GECI (f1-3). The 10-20 minutes resting state session of simultaneous fMRI/WOI were preprocessed and filtered to <0.1Hz and segmented to 400s with 50% overlap before performing a cross-correlation analysis. The average results were exhibited in color plots on various comparisons between hemodynamic signals (EPI, ZTE, HbT) and fluorescence (raw or corrected) in GEVI (f1) and GECI (f2 and f3).

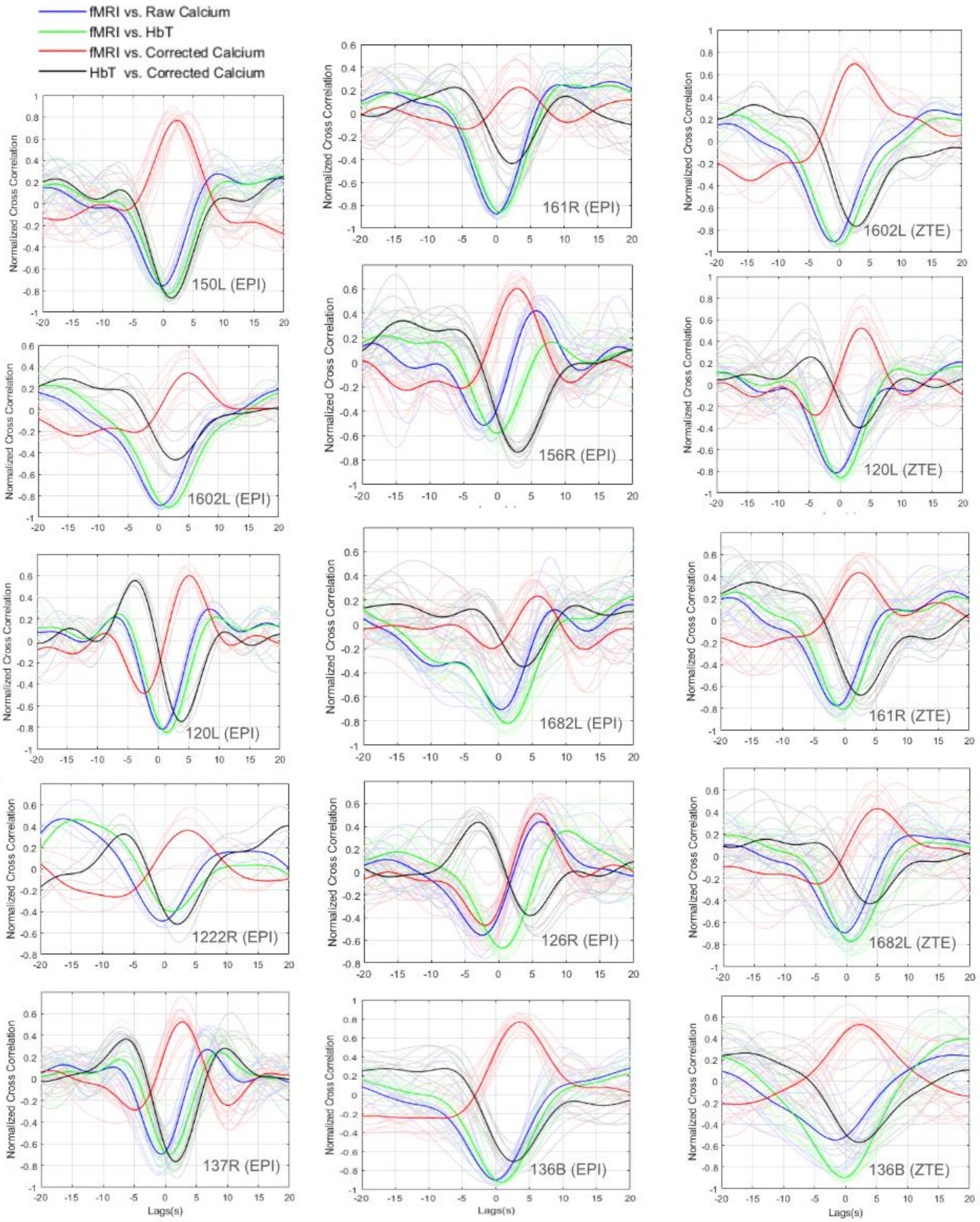


**Fig. S5** Individual cross-correlation data between fMRI and WOI as an example of the barrel-field network in the resting state in GCaMP mice (n=10 EPI scans, 5/10 with additional ZTE scans). Each scan lasted for 10-20 min. The cross correlation was calculated on <0.1 Hz signals, averaged from multiple segments of 200s long and 50% overlap for each scan.  As expected, all the data exhibited a high correlation between the fMRI and HbT absorption signals. The raw calcium fluorescence signals were largely similar to the HbT absorption signals, with a negative correlation with fMRI and almost no time lag. After HbT regression, the corrected calcium signal was purified and positively correlated with fMRI data with time lags of a few seconds.

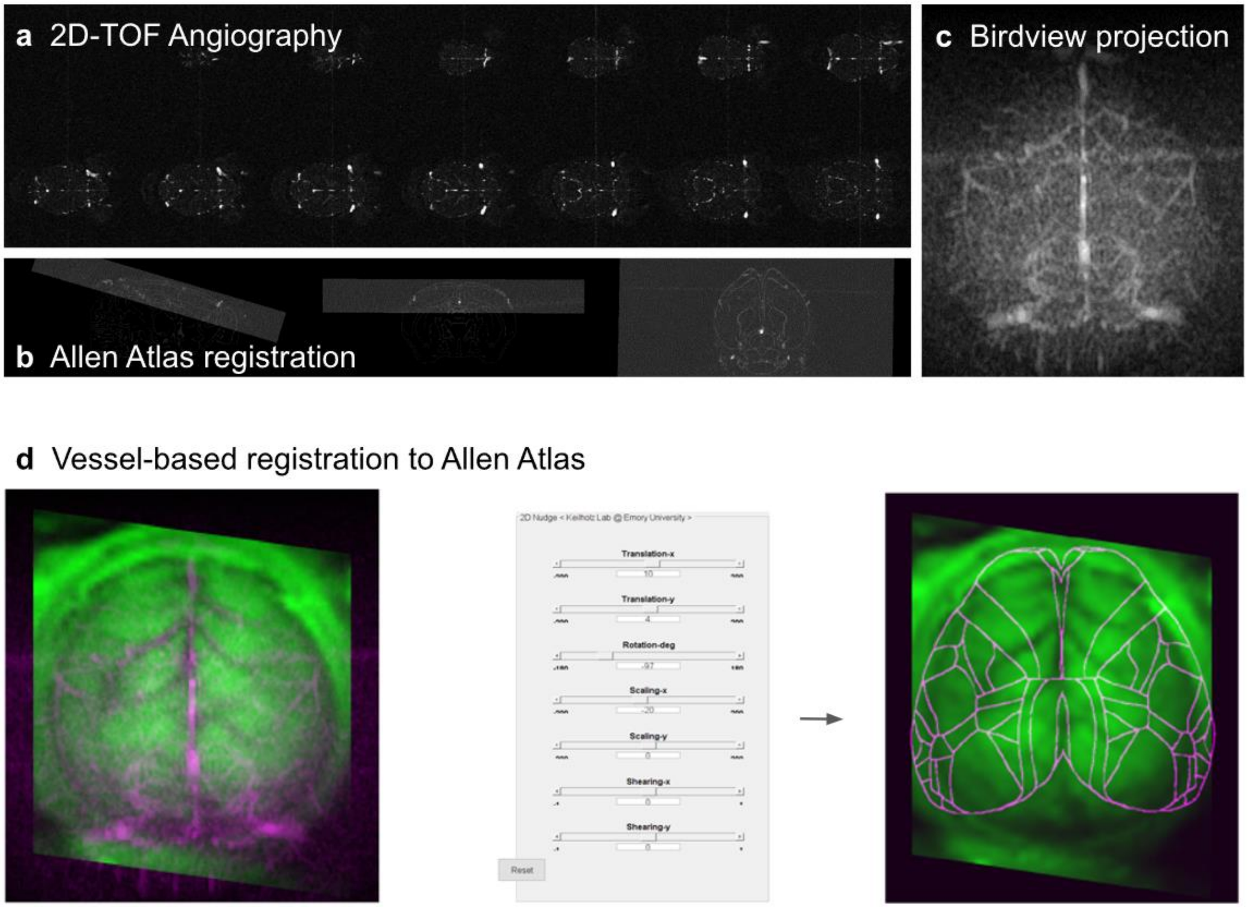


**Fig.S6** WOI image atlas registration method. (a) Cortical 3D vessels were obtained by MRI 2D-TOF (14 slices, 0.4 mm thickness). (b) Angiography images were initially registered into the Allen 3D atlas space using affine transformation and averaged into a 2D cortical vessel image (c). Spatially normalized vessels (d) were used for the WOI image registration in the Allen cortical atlas.

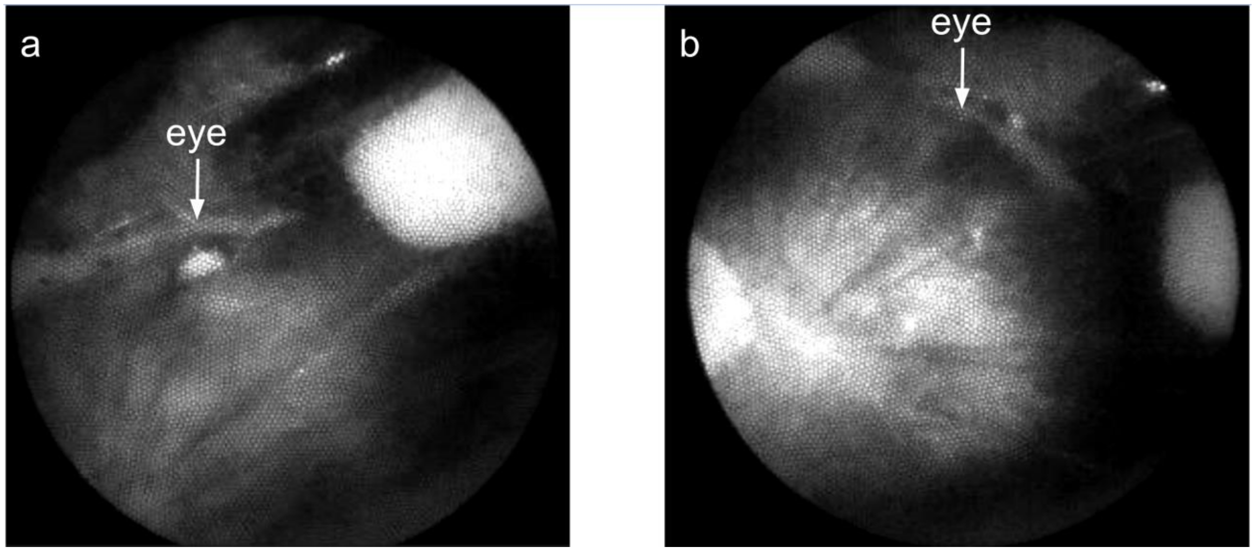


Fig. S7 Pupil comparison between conditions of isoflurane anesthesia and awake state. In our pilot studies, we observed clear pupil reflection bright light during awake state (a) but the pupil reflection was rarely seen during isoflurane anesthesia (b). During simultaneous WOI, lighting on brain may also be another possible reason that stimulated retina from back and caused pupil shrink or eyeball turn in reflection. Therefore, the face cam was used for monitoring whisker behavior for estimating anesthesia depth only.

**Supplementary Table S1:** conventional fMRI and simultaneous fMRI/WOI comparison.

| | **Conventional fMRI** | **fMRI with WOI** |
|---|---|---|
| Non-brain tissue (overhead) | Intact | Removed |
| Surface coil | Flat | Curve-fitted to head |
| Coil-brain distance | Variable | Minimum |
| Coil positioning | Variable | Highly reproducible |
| Head fixation | Ear bars (on skin) | Implanted headpiece (on bone) |

**Supplementary Table S2:** the WOI trade-off profile between GCaMP6f and JEDI

| **Feature** [1–6] | **GCaMP6f (Calcium)** | **JEDI-1P (Voltage)** |
|---|---|---|
| **Physical Localization** | Cytoplasmic (fills the entire cell body) | Membrane-bound (restricted to thin outer shell) |
| **Kinetics (On / Off)** | ~50 ms / ~200–300 ms | **~1.6 ms / ~1.6 ms** |
| **Maximum Frequency** | ~4–5 Hz | **> 60–100 Hz** |
| **Sensitivity (Δ F/F□)** | Extremely high (> 100% per action potential burst) | **Moderate to High (~55% maximum dynamic range)** |
| **Baseline Photon Budget** | Fills cell volume; requires modest illumination | Restricted to membrane; requires ultra-high illumination |
| **Photobleaching Rate** | Negligible over hours of imaging | Low for a GEVI, but still limits trial lengths |
| **Tissue Depth Limit** | Surface layers (~100–200 µm with 1P widefield) | Strictly surface layers (highly vulnerable to 1P light scattering) |

**Supplementary Table S3:** comparison with fiber bundle methods

| Feature | Lake et al[7]. | Chen et al[8]. | This work |
|---|---|---|---|
| Optical transmission | Fiber bundle (SCHOTT, ~40%) | Fiber bundle (Zibra Corporation, not specified) | Tube-lens (>98%) |
| Transparent skull rebuilt | No | No | Yes |
| Objective lens distance/NA | Short/Large | Short/Large | Long/Small |
| Illumination angle | 90° | 90° | 15° |
| fMRI sequences | EPI only | EPI only | EPI + ZTE |
| Fluorescence range | Whole brain | Local cortical regions | Whole brain |
| Voltage indicator | No | No | Yes (JEDI) |
| Chronic window duration | Not specified | Up to 4 months | Up to 15 months |
| Hemodynamic correction | Yes (violet regression) | No | Yes (HbT regression) |

References

1. Nguyen TN, Shalaby RA, Lee E, et al. Ultrafast optical imaging techniques for exploring rapid neuronal dynamics. *Neurophotonics*. 2025;12(S1):S14608. doi:10.1117/1.NPh.12.S1.S14608

2. Penzkofer A. Voltage sensitive probes for membrane potential determination in life science. *Res*. 2026;7:101497. doi:10.1016/j.nexres.2026.101497

3. Lu X, Wang Y, Liu Z, Gou Y, Jaeger D, St-Pierre F. Widefield imaging of rapid pan-cortical voltage dynamics with an indicator evolved for one-photon microscopy. *Nat Commun*. 2023;14:6423. doi:10.1038/s41467-023-41975-3

4. Yang S, McDonald AJ, Lu X, et al. A red-emitting, genetically encoded indicator for two-photon voltage recording in vivo. *bioRxiv: The Preprint Server for Biology*. Preprint posted online June 3, 2026:2026.06.01.726307. doi:10.64898/2026.06.01.726307

5. Lu X, Wang Y, Liu Z, Gou Y, Jaeger D, St-Pierre F. Detecting rapid pan-cortical voltage dynamics in vivo with a brighter and faster voltage indicator. *bioRxiv*. Preprint posted online August 31, 2022:2022.08.29.505018. doi:10.1101/2022.08.29.505018

6. Nikolaev DM, Metelkina EM, Shtyrov AA, Li F, Panov MS, Ryazantsev MN. Noise Sources and Strategies for Signal Quality Improvement in Biological Imaging: A Review Focused on Calcium and Cell Membrane Voltage Imaging. *Biosensors*. 2026;16(1):31. doi:10.3390/bios16010031

7. Lake EM, Ge X, Shen X, et al. Simultaneous cortex-wide fluorescence Ca2+ imaging and whole-brain fMRI. *Nat Methods*. 2020;17(12):1262-1271. doi:10.1038/s41592-020-00984-6

8. Chen Z, Chen Y, Gezginer I, et al. Non-invasive large-scale imaging of concurrent neuronal, astrocytic, and hemodynamic activity with hybrid multiplexed fluorescence and magnetic resonance imaging (HyFMRI). *Light Sci Appl*. 2025;14(1):341. doi:10.1038/s41377-025-02003-9
